# Dithiothreitol causes toxicity in *C. elegans* by modulating the methionine-homocysteine cycle

**DOI:** 10.1101/2021.11.16.468906

**Authors:** Gokul G, Jogender Singh

## Abstract

The redox reagent dithiothreitol (DTT) causes stress in the endoplasmic reticulum (ER) by disrupting its oxidative protein folding environment, which results in the accumulation and misfolding of the newly synthesized proteins. DTT may potentially impact cellular physiology by ER-independent mechanisms; however, such mechanisms remain poorly characterized. Using the nematode model *Caenorhabditis elegans*, here we show that DTT toxicity is modulated by the bacterial diet. Specifically, the dietary component vitamin B12 alleviates DTT toxicity in a methionine synthase-dependent manner. Using a forward genetic screen, we discover that loss-of-function of *R08E5*.*3*, an S-adenosylmethionine (SAM)-dependent methyltransferase, confers DTT resistance. DTT upregulates *R08E5*.*3* expression and modulates the activity of the methionine-homocysteine cycle. Employing genetic studies, we establish that DTT toxicity is a result of the depletion of SAM. Finally, we show that a functional IRE-1/XBP-1 unfolded protein response pathway is required to counteract toxicity at high, but not low, DTT concentrations.

## INTRODUCTION

Diverse biological processes occur via oxidation-reduction (redox) reactions, and therefore, a redox balance is essential for cellular and organismal homeostasis ^1^. A consequence of biological redox reactions is the generation of reactive oxygen species (ROS). The physiological flux of ROS regulates cellular processes essential for cell survival, maintenance, and aging ^1,2^. In contrast, excessive ROS leads to oxidative stress, a hallmark of many pathological states ^2^. To combat excessive ROS, the cellular systems have developed antioxidant networks, including thioredoxin, glutathione, glutathione peroxidases, catalase, and superoxide dismutases ^3^. In addition to endogenous antioxidants, dietary antioxidant supplements are taken under healthy as well as pathological conditions to counteract ROS ^3^. Animal and cell model studies suggest that thiol-based antioxidants, including glutathione, N-acetylcysteine (NAC), and dithiothreitol (DTT), may attenuate ROS-induced pathological conditions such as hepatic, hematopoietic, intestinal, and renal injuries ^4–6^, Alzheimer’s disease ^7^, and dopamine-induced cell death ^8^. While the thiol-based antioxidants appear to improve various pathological conditions, their broad effects on cellular physiology remain poorly characterized. Recently, it was shown that glutathione and NAC could have detrimental effects on the health and lifespan of *C. elegans* ^9^. Therefore, a better characterization of the physiological effects of various thiol antioxidants is necessary to harness their therapeutic value.

DTT is a commonly used redox reagent that contains two thiol groups ^10^ and is known to reduce protein disulfide bonds ^11^. Because the endoplasmic reticulum (ER) has an oxidative environment conducive to disulfide bond formation, DTT reduces disulfide bonds in the ER ^12^. Therefore, DTT is widely used as an ER-specific stressor ^12,13^. While DTT has been shown to improve several ROS-based pathological conditions ^4–6,8^, high amounts of DTT are toxic and lead to cell death which is thought to be mediated by increased ER stress ^14–16^. However, some studies suggest that DTT may induce apoptosis by generating hydrogen peroxide and oxidative stress ^17,18^. Similarly, several other DTT-mediated phenotypes are shown to be independent of ER stress ^19–22^. Therefore, how DTT impacts cellular physiology remains to be fully understood.

Here, using the *C. elegans* model, we characterized the physiological effects of DTT. We showed that DTT causes developmental defects in *C. elegans* in a bacterial diet-dependent manner. The dietary component vitamin B12 alleviated development defects caused by DTT by acting as a cofactor for methionine synthase. To understand the interplay between DTT and vitamin B12, we isolated *C. elegans* mutants in a forward genetic screen that were resistant to DTT toxicity. Loss-of-function of *R08E5*.*3*, a SAM-dependent methyltransferase, imparted DTT resistance. DTT resulted in the upregulation of *R08E5*.*3* expression and, therefore, modulated the methionine-homocysteine cycle. DTT resulted in the depletion of SAM, and supplementation of methionine and choline could rescue DTT toxicity. Modulation of the methionine-homocysteine cycle by DTT also resulted in the upregulation of the ER and mitochondrial unfolded protein response (UPR) pathways. Finally, we showed that while DTT toxicity primarily occurred via the modulation of the methionine-homocysteine cycle, a functional IRE-1/XBP-1 UPR pathway was required to counteract DTT toxicity.

## RESULTS

### DTT affects *C. elegans* development in a bacterial diet-dependent manner

DTT is known to be toxic to *C. elegans* and affects its development ^23^. We first studied *C. elegans* development retardation at different concentrations of DTT. In the presence of *E. coli* OP50 diet, *C. elegans* was at the L1 or L2 stage at 5 mM and higher DTT concentrations at 72 hours of hatching, while the control animals without DTT had developed to adulthood at the same time (Figures 1A and 1B). We also studied development retardation in the presence of DTT on *E. coli* HT115 bacterium, a strain commonly used for feeding-based RNA interference (RNAi). Surprisingly, we observed that *C. elegans* developed much better on DTT in the presence of *E. coli* HT115 diet than *E. coli* OP50 diet (Figures 1A-1C). These results suggested that some bacterial components might be able to modulate DTT toxicity.

**Figure 1.**
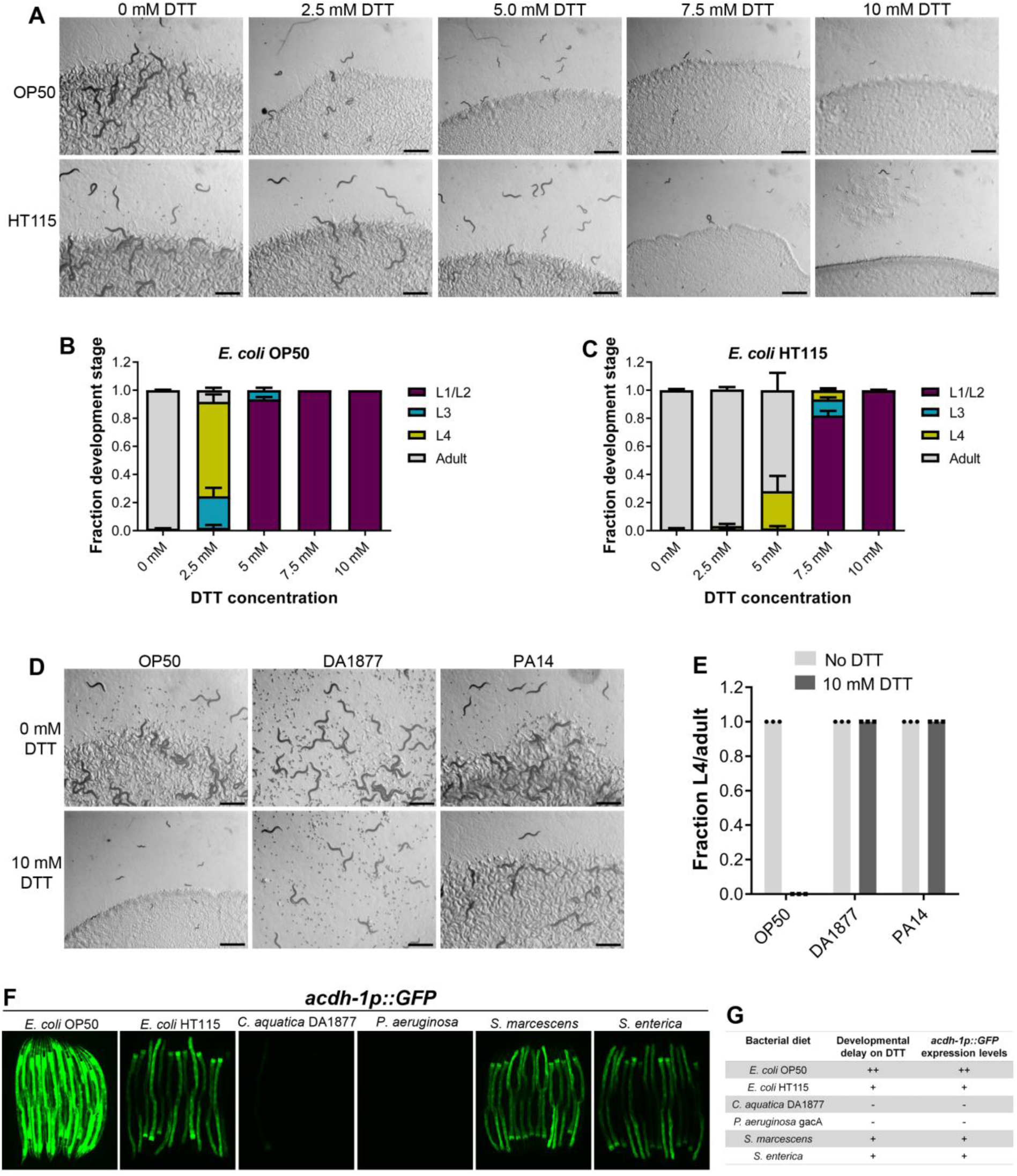
DTT affects *C. elegans* development in a diet-dependent manner. (A) Representative images of wild-type N2 *C. elegans* on various concentrations of DTT on *E. coli* OP50 and *E. coli* HT115 diets after 72 hours of hatching at 20°C. Scale bar = 1 mm. (B and C) Quantification of different developmental stages of wild-type N2 *C. elegans* on various concentrations of DTT on *E. coli* OP50 (B) and *E. coli* HT115 (C) diets after 72 hours of hatching at 20°C (n = 3 biological replicates; animals per condition per replicate > 80). (D) Representative images of wild-type N2 *C. elegans* after 72 hours of hatching at 20°C on(E) *E. coli* OP50, *C. aquatica* DA1877, and *P. aeruginosa* PA14 gacA mutant diets containing either 0 mM or 10 mM DTT. Scale bar = 1 mm. (E) Fraction L4 or adult wild-type N2 *C. elegans* after 72 hours of hatching at 20°C on *E. coli* OP50, *C. aquatica* DA1877, and *P. aeruginosa* PA14 gacA mutant diets containing either 0 mM or 10 mM DTT (n = 3 biological replicates; animals per condition per replicate > 80). (F) Representative fluorescence images of *acdh-1p::GFP* animals grown on various bacterial diets. (G) Table summarizing the effects of bacterial diet on DTT-induced developmental delay and on GFP levels in *acdh-1p::GFP* animals.

To gain an understanding of the potential bacterial components that might modulate DTT toxicity, we studied the development of *C. elegans* in the presence of DTT on different bacterial diets, including *Comamonas aquatica* DA1877, *Pseudomonas aeruginosa* PA14 *gacA* mutant, *Salmonella enterica*, and *Serratia marcescens*. Because *P. aeruginosa* virulence might affect the development of *C. elegans*, we used the *gacA* mutant, which has drastically reduced virulence ^24^. Interestingly, *C. elegans* showed no developmental defects on *C. aquatica* DA1877 and *P. aeruginosa* PA14 diets in the presence of 10 mM DTT (Figures 1D and 1E). Similarly, *C. elegans* developed much better on *S. enterica* and *S. marcescens* diets compared to the *E. coli* OP50 diet at lower DTT concentrations (Figures S1A-S1D). These results showed that bacterial diet has a major effect on DTT toxicity, and likely some bacterial diet component(s) modulated DTT toxicity.

Previous studies have shown that relative to the *E. coli* OP50 diet, a *C. aquatica* DA1877 diet accelerates *C. elegans* development primarily because it contains higher levels of vitamin B12 ^25^. Similarly, *E. coli* HT115 and *P. aeruginosa* PA14 diets are known to have higher amounts of vitamin B12 than the *E. coli* OP50 diet ^25,26^. The amount of dietary vitamin B12 available to *C. elegans* can be reliably determined by a dietary sensor strain expressing green fluorescent protein (GFP) under the promoter of acyl-coenzyme A dehydrogenase (*acdh-1*) gene ^25^. When the animals experience low levels of vitamin B12, *acdh-1p::GFP* is strongly induced, and the level of GFP expression reduces as the animals obtain increasing levels of vitamin B12. Using *acdh-1p::GFP* animals, we first tested the levels of vitamin B12 in the animals on different bacterial diets compared to the *E. coli* OP50 diet. As shown in Figure 1F, the animals show very bright GFP expression on the *E. coli* OP50 diet, while the GFP levels were diminished on *E. coli* HT115, *S. enterica*, and *S. marcescens* diets. On the other hand, the animals showed no GFP expression on *C. aquatica* DA1877 and *P. aeruginosa* PA14 diets. The levels of GFP expression in *acdh-1p::GFP* animals on different bacterial diets correlated perfectly with the developmental defects on these bacterial diets in the presence of DTT (Figure 1G).

### Vitamin B12 alleviates DTT toxicity via methionine synthase

The correlation between the GFP levels in *acdh-1p::GFP* animals and the developmental defects of animals in the presence of DTT on different bacterial diets suggested that vitamin B12 could be the microbial component that alleviates DTT toxicity in *C. elegans*. To test this, we studied the development of N2 animals on *E. coli* OP50 diet containing 10 mM DTT and supplemented with 50 nM vitamin B12. In contrast to the development retardation on *E. coli* OP50 diet containing 10 mM DTT, the animals developed to adulthood on *E. coli* OP50 diet containing 10 mM DTT supplemented with 50 nM vitamin B12 (Figures 2A and 2B). We further asked whether the DTT-mediated development retardation could be reversed by supplementation of vitamin B12. To this end, vitamin B12 was added to the L1-arrested animals after 72 hours of hatching on *E. coli* OP50 plates containing 10 mM DTT. After 48 hours of incubation post vitamin B12 supplementation, more than 95% of the animals developed to adulthood (Figures S2A and S2B). On the other hand, the control animals on *E. coli* OP50 plates containing 10 mM DTT, but without vitamin B12 supplementation, were still at the L1-L2 stages after 120 hours of hatching. Taken together, these studies showed that vitamin B12 is sufficient to alleviate and revert the toxic effects of DTT.

**Figure 2.**
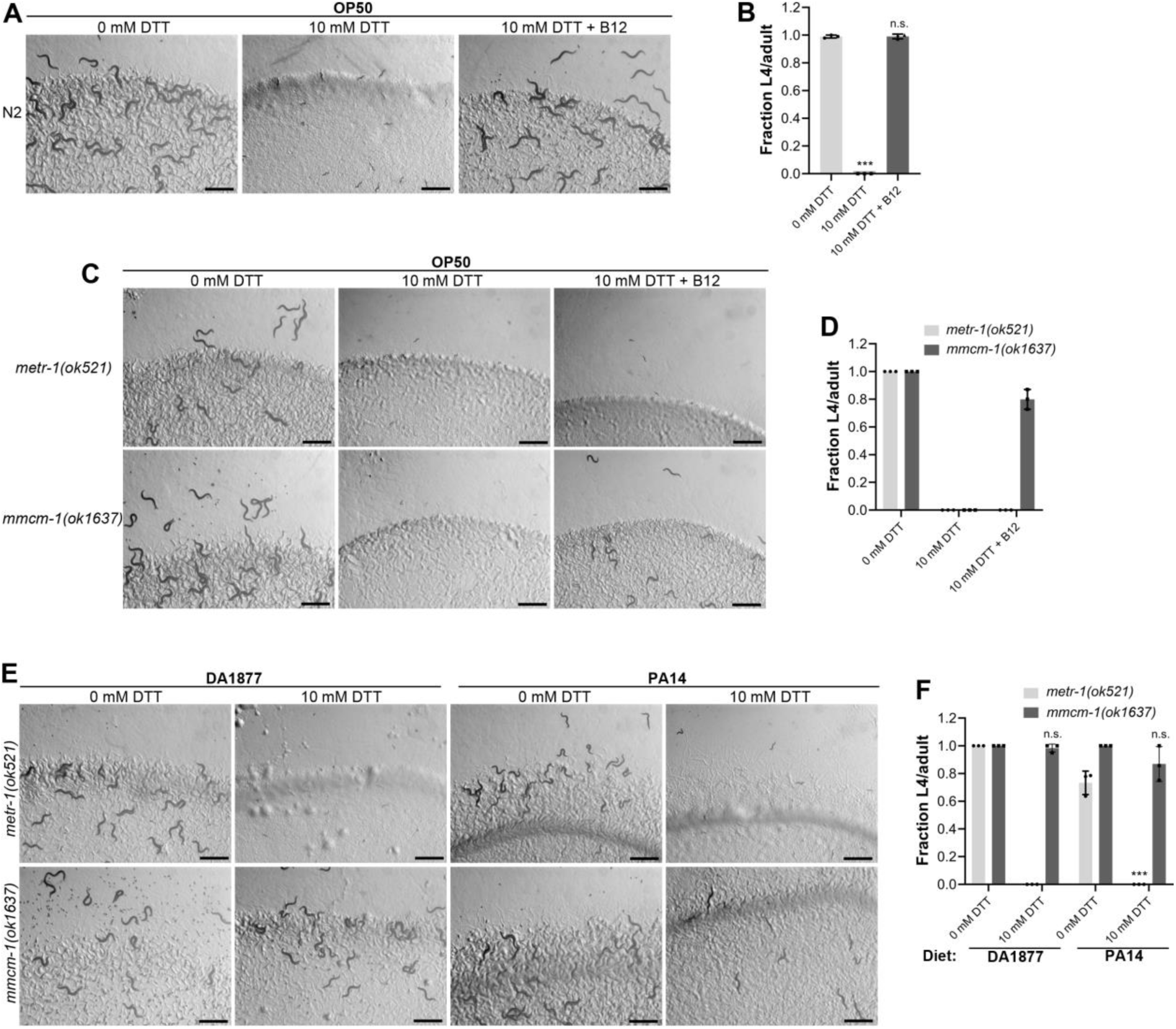
Vitamin B12 alleviates DTT toxicity via methionine synthase. (A) Representative images of wild-type N2 *C. elegans* after 72 hours of hatching at 20°C on(B) *E. coli* OP50 diet containing 0 mM DTT, 10 mM DTT, and 10 mM DTT supplemented with 50 nM vitamin B12. Scale bar = 1 mm. (B) Fraction L4 or adult wild-type N2 *C. elegans* after 72 hours of hatching at 20°C on *E. coli* OP50 diet containing 0 mM DTT, 10 mM DTT, and 10 mM DTT supplemented with 50 nM vitamin B12. p values are relative to 0 mM DTT. ***p < 0.001 via the t test. n.s., non-significant (n = 3 biological replicates; animals per condition per replicate > 80). (C) Representative images of *metr-1(ok521)* and *mmcm-1(ok1637)* animals after 72 hours of hatching at 20°C on *E. coli* OP50 diet containing 0 mM DTT, 10 mM DTT, and 10 mM DTT supplemented with 50 nM vitamin B12. Scale bar = 1 mm. (D) Fraction L4 or adult *metr-1(ok521)* and *mmcm-1(ok1637)* animals after 72 hours of hatching at 20°C on *E. coli* OP50 diet containing 0 mM DTT, 10 mM DTT, and 10 mM DTT supplemented with 50 nM vitamin B12 (n = 3 biological replicates; animals per condition per replicate > 60). (E) Representative images of *metr*-1(ok521) and *mmcm*-1(ok1637) animals after 72 hours of hatching at 20°C on *C. aquatica* DA1877 and *P. aeruginosa* PA14 *gacA* mutant diets containing either 0 mM or 10 mM DTT. Scale bar = 1 mm. (F) Fraction L4 or adult *metr*-1(ok521) and *mmcm*-1(ok1637) animals after 72 hours of hatching at 20°C on *C. aquatica* DA1877 and *P. aeruginosa* PA14 *gacA* mutant diets containing either 0 mM or 10 mM DTT. p values are relative to the corresponding 0 mM DTT condition for each mutant. ***p < 0.001 via the t test. n.s., non-significant (n = 3 biological replicates; animals per condition per replicate > 60).

Vitamin B12 is an essential cofactor for two metabolic enzymes: methylmalonyl-CoA mutase (MMCM-1 in *C. elegans*), which converts methylmalonyl-CoA to succinyl-CoA in the propionyl-CoA breakdown pathway, and methionine synthase (METR-1 in *C. elegans*), which converts homocysteine to methionine in the methionine-homocysteine cycle. We asked whether one or both of these enzymes are required for alleviating DTT toxicity. While vitamin B12 supplementation rescued the toxic effects of DTT in *mmcm-1(ok1637)* animals, it failed to do so in *metr-1(ok521)* animals (Figures 2C and 2D). The *metr-1(ok521)* animals were at the L1 stage on *E. coli* OP50 diet containing 10 mM DTT regardless of vitamin B12 supplementation (Figure 2C). Further, *mmcm-1(ok1637)* animals did not show any significant development retardation effects when grown on *C. aquatica* DA1877 and *P. aeruginosa* PA14 diets containing 10 mM DTT (Figures 2E and 2F). On the other hand, *metr-1(ok521)* animals failed to develop when grown on *C. aquatica* DA1877 and *P. aeruginosa* PA14 diets containing 10 mM DTT (Figures 2E and 2F). Taken together, these studies showed that vitamin B12 could alleviate DTT toxicity via the methionine synthase enzyme.

### DTT toxicity is mediated via an S-adenosylmethionine (SAM)-dependent methyltransferase

To understand why DTT toxicity depended on vitamin B12 and the methionine-homocysteine cycle, we carried out a forward genetic screen for mutants that would develop to L4/adult stage at 72 hours post-hatching on *E. coli* OP50 diet containing 10 mM DTT. From a screen of approximately 50,000 ethyl methanesulfonate (EMS)-mutagenized haploid genomes, twelve mutants were isolated that exhibited DTT resistance and developed on *E. coli* OP50 diet containing 10 mM DTT (Figures S3A and S3B). To identify the causative mutations, we conducted the whole-genome sequencing of all the twelve mutants after backcrossing them six times with the wild-type N2 strain. The sequenced genomes were aligned with the reference genome of *C. elegans*. After subtraction of the common variants, linkage maps of single nucleotide polymorphisms (SNPs) were obtained (Figure S4). All of the mutants had linkage to chromosome V. Analysis of the protein-coding genes carrying mutations in the linked regions of each mutant revealed that all of the mutants had a mutation in the gene *R08E5*.*3* (Figures 3A and 3B), which we named as *drm-1* (***D***TT-***r***esistant ***m***utants). *drm-1* is predicted to encode a protein with a SAM-dependent methyltransferase domain. SAM-dependent methyltransferases catalyze diverse methylation reactions where a methyl group is transferred to the substrate from SAM, resulting in the conversion of SAM to S-adenosylhomocysteine (SAH) in the methionine-homocysteine cycle. Fortuitously, the allele *gk231506* available in the Caenorhabditis Genetics Center (CGC) has the same molecular change in the protein DRM-1 as the allele *jsn6* obtained in our screen has. *drm-1(gk231506)* animals showed DTT resistance and developed on *E. coli* OP50 diet containing 10 mM DTT (Figures 3C and 3D).

**Figure 3.**
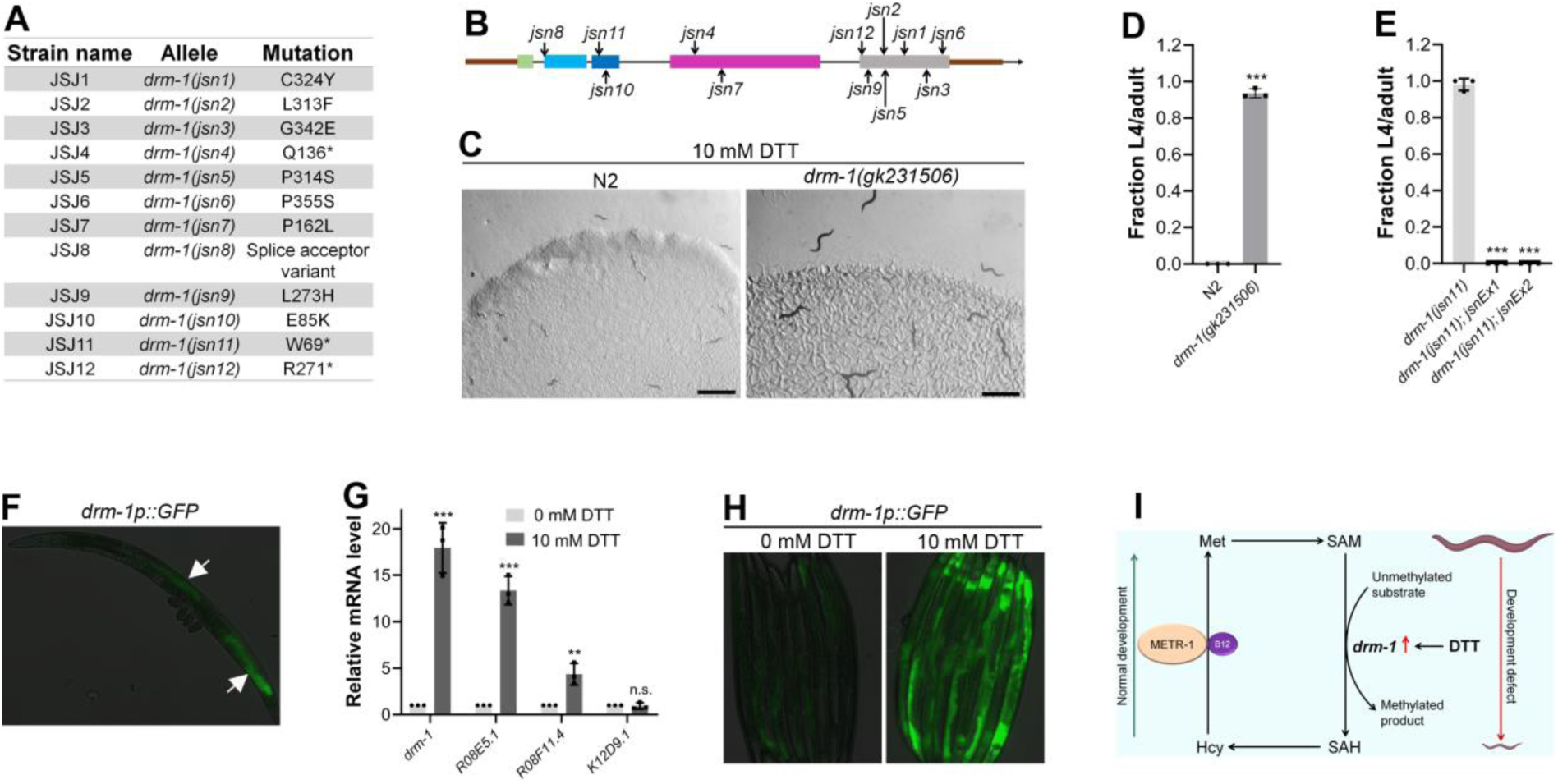
DTT causes developmental toxicity via *drm-1*. (A) Table summarizing the alleles of *drm-1* identified by whole-genome sequencing in different DTT-resistant strains. The corresponding amino acid changes in the DRM-1 protein are also shown. (B) Mapping of the *drm-1* alleles identified in the forward genetic screen. (C) Representative images of N2 and *drm-1(gk231506)* animals after 84 hours of hatching at 20°C on *E. coli* OP50 diet containing 10 mM DTT. Scale bar = 1 mm. (D) Fraction L4 or adult N2 and *drm-1(gk231506)* animals after 84 hours of hatching at 20°C on *E. coli* OP50 diet containing 10 mM DTT. ***p < 0.001 via the t test (n = 3 biological replicates; animals per condition per replicate > 50). (E) Fraction L4 or adult *drm-1(jsn11), drm-1(jsn11);jsnEx1*, and *drm-1(jsn11);jsnEx2* animals after 72 hours of hatching at 20°C on *E. coli* OP50 diet containing 10 mM DTT. *jsnEx1* and *jsnEx2* represent two independent extrachromosomal arrays containing *drm-1p::drm-1::SL2::GFP* and *myo-2p::mCherry*. ***p < 0.001 via the t test (n = 3 biological replicates; animals per condition per replicate > 50). (F) Representative fluorescence image of *drm-1p::GFP* animal. The white arrows point at the intestinal regions showing GFP expression. (G) Gene expression analysis of N2 animals grown on *E. coli* OP50 diet containing 0 mM DTT until the young adult stage, followed by incubation on *E. coli* OP50 diet containing 0 mM or 10 mM DTT for 4 hours. ***p < 0.001 and **p < 0.01 via the t test. n.s., non-significant (n = 3 biological replicates). (H) Representative fluorescence images of *drm-1p::GFP* animals grown on *E. coli* OP50 diet containing 0 mM DTT until the young adult stage, followed by incubation on *E. coli* OP50 diet containing 0 mM or 10 mM DTT for 8 hours. (I) Model depicting the effects of DTT and vitamin B12 on *C. elegans* development via the methionine-homocysteine cycle.

To confirm that resistance to DTT toxicity is imparted by loss-of-function mutations in *drm-1*, we created *drm-1(jsn11)* animals expressing *drm-1* under its own promoter. As shown in Figure 3E, the DTT-resistant phenotype of *drm-1(jsn11)* animals was completely rescued in these animals. We further determined the expression pattern of *drm-1* by expressing GFP driven by its own 918-bp promoter in wild-type N2 animals. As shown in Figure 3F, *drm-1* is expressed in the intestine.

### DTT modulates the methionine-homocysteine cycle by upregulating *drm-1* expression

We next asked how DTT exerted its toxic effects via *drm-1*. In the fungus *Aspergillus niger*, DTT is known to highly upregulate two SAM-dependent methyltransferases ^20^. We asked whether DTT also upregulated the SAM-dependent methyltransferase *drm-1*, and any other closely related methyltransferases. *drm-1* shares high sequence similarity with three other SAM-dependent methyltransferases in *C. elegans, R08E5*.*1* (BLAST e value: 1e-168), *R08F11*.*4* (BLAST e value: 1e-166), and *K12D9*.*1* (BLAST e value: 2e-152). Quantitative reverse transcription-PCR (qRT-PCR) analysis showed that DTT resulted in significant upregulation of the transcript levels of *drm-1, R08E5*.*1*, and *R08F11*.*4* (Figure 3G). The enhanced expression of *drm-1* was confirmed in a reporter strain expressing GFP under the promoter of *drm-1* (Figure 3H). Taken together, these results suggested that by increasing expression of *drm-1*, DTT modulates the methionine-homocysteine cycle resulting in toxic effects that can be reversed by supplementation of vitamin B12 (Figure 3I).

Modulation of the methionine-homocysteine cycle or low dietary vitamin B12 are known to upregulate the mitochondrial UPR ^27,28^. Since DTT modulates the methionine-homocysteine cycle and requires vitamin B12 to reverse those effects, we asked whether DTT can also lead to the upregulation of the mitochondrial UPR. To this end, we exposed the adult *hsp-6p::GFP* animals to DTT. Compared to the control animals, animals exposed to 10 mM DTT had upregulation of the mitochondrial UPR (Figure S5). Supplementation of vitamin B12 rescued the upregulation of mitochondrial UPR by DTT, indicating that DTT induced mitochondrial UPR via the methionine-homocysteine cycle.

### DTT toxicity is a result of SAM depletion

Next, we asked how perturbations of the methionine-homocysteine cycle activity led to DTT toxicity. Since DTT exerts its toxicity via the SAM-dependent methyltransferase DRM-1, we postulated that either depletion of the metabolites upstream of DRM-1 activity (methionine and SAM) or accumulation of metabolites downstream of DRM-1 activity (SAH and homocysteine) or a combination of both would result in DTT toxicity. To discriminate between these possibilities, we carried out methionine supplementation assays in the presence of DTT. We reasoned that if DTT toxicity is because of methionine and/or SAM depletion, supplementation of methionine should attenuate DTT toxicity. On the other hand, if DTT toxicity is caused by the accumulation of SAH and/or homocysteine, supplementation of methionine should enhance DTT toxicity as methionine supplementation would likely enhance levels of SAH and/or homocysteine. The wild-type N2 animals had improved development when 10 mM DTT containing *E. coli* OP50 diet was supplemented with varying concentrations of methionine (Figures 4A and 4B). Methionine supplementation alleviated DTT toxicity in a dose-dependent manner (Figures 4A and 4B). We observed that *metr-1(ok521)* animals that failed to develop upon vitamin B12 supplementation in the presence of DTT (Figure 2C and 2D), developed upon supplementation of methionine (Figures 4A and 4C). These results suggested that depletion of methionine and/or SAM is the primary cause of DTT toxicity.

**Figure 4.**
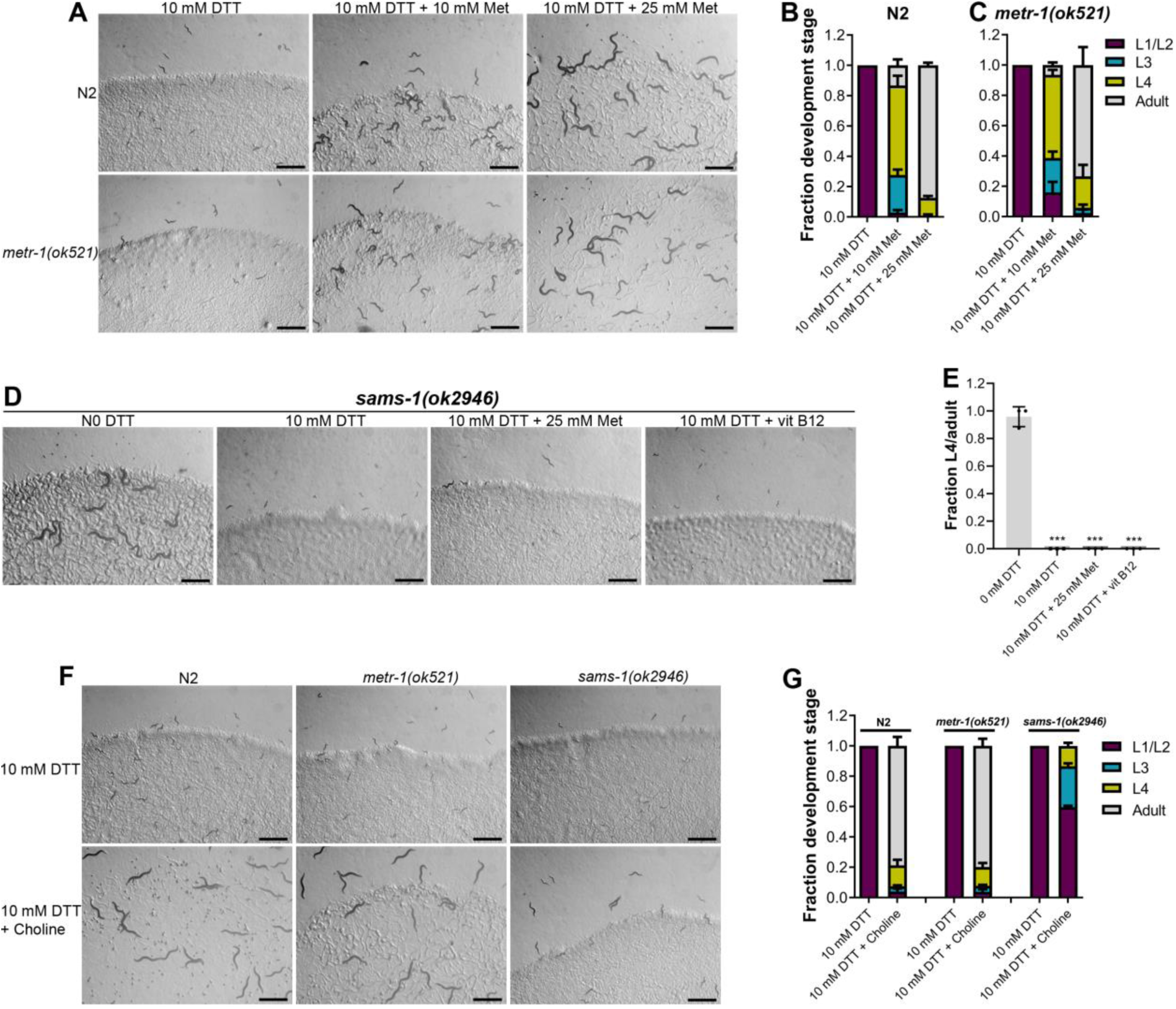
Methionine and choline supplementation alleviate DTT toxicity. (A) Representative images of N2 and *metr-1(ok521)* animals after 5 days of hatching at 20°C on *E. coli* OP50 diet containing 10 mM DTT, 10 mM DTT supplemented with 10 mM methionine, and 10 mM DTT supplemented with 25 mM methionine. Scale bar = 1 mm. (B and C) Quantification of different developmental stages of N2 (B) and *metr-1(ok521)* (C) animals after 5 days of hatching at 20°C on *E. coli* OP50 diet containing 10 mM DTT, 10 mM DTT supplemented with 10 mM methionine, and 10 mM DTT supplemented with 25 mM methionine (n = 3 biological replicates; animals per condition per replicate > 60). (D) Representative images of *sams-1(ok2946)* animals grown on *E. coli* OP50 diet containing 0 mM DTT, 10 mM DTT, 10 mM DTT supplemented with 25 mM methionine, and 10 mM DTT supplemented with 50 nM vitamin B12. The animals were grown for 5 days on the plates containing methionine and for 3 days under all other conditions. Scale bar = 1 mm. (E) Fraction L4 or adult *sams-1(ok2946)* animals grown on *E. coli* OP50 diet containing 0 mM DTT, 10 mM DTT, 10 mM DTT supplemented with 25 mM methionine, and 10 mM DTT supplemented with 50 nM vitamin B12. The animals were grown for 5 days on the plates containing methionine and for 3 days under all other conditions. ***p < 0.001 via the t test (n = 3 biological replicates; animals per condition per replicate > 50). (F) Representative images of N2, *metr-1(ok521)*, and *sams-1(ok2946)* animals after 4 days of hatching at 20°C on *E. coli* OP50 diet containing 10 mM DTT and 10 mM DTT supplemented with 80 mM choline. Scale bar = 1 mm. (G) Quantification of different developmental stages of N2, *metr-1(ok521)*, and *sams-1(ok2946)* animals after 4 days of hatching at 20°C on *E. coli* OP50 diet containing 10 mM DTT and 10 mM DTT supplemented with 80 mM choline (n = 3 biological replicates; animals per condition per replicate > 50).

Next, we studied whether DTT toxicity was an outcome of depletion of methionine per se or its downstream metabolite SAM. The first step in the methionine-homocysteine cycle involves the conversion of methionine into SAM by methionine adenosyltransferase (S-adenosylmethionine synthetase or SAMS in *C. elegans*). Animals having knockdown of *sams-1* have defects in methionine to SAM conversion and exhibit significantly reduced levels of SAM ^29^. Because *sams-1* mutants have reduced conversion of methionine to SAM, supplementation of methionine would alleviate DTT toxicity in *sams-1* mutants only if DTT toxicity is caused by depletion of methionine and not SAM. Supplementation of methionine did not improve the development of *sams-1(ok2946)* animals on *E. coli* OP50 diet containing 10 mM DTT (Figures 4D and 4E). Further, *sams-1(ok2946)* animals also failed to develop on 10 mM DTT containing *E. coli* OP50 diet supplemented with 50 nM vitamin B12 (Figures 4D and 4E). Taken together, these results suggested that SAM depletion is the primary cause of DTT toxicity.

SAM is the universal methyl donor in the cell and regulates a vast array of cellular activities, including the synthesis of phosphatidylcholine by methylation of phosphoethanolamine ^30^. Several of the SAM functions, including lipogenesis ^29^, gene expression changes ^31^, innate immunity ^32^, are regulated by phosphatidylcholine. Phosphatidylcholine can also be synthesized from choline by an alternate pathway independent of SAM ^30^. Importantly, the SAM-dependent phenotypes that require phosphatidylcholine can be rescued by supplementation of choline ^29,31,32^. We asked whether choline could also attenuate DTT toxicity. Supplementation of choline rescued the developmental defects in N2 and *metr-1(ok521)* animals on an *E. coli* OP50 diet containing 10 mM DTT (Figures 4F and 4G). The rescue of DTT toxicity was only partial in *sams-1(ok2946)* animals (Figures 4F and 4G). These results suggested that phosphatidylcholine is a major, but not the sole, SAM product responsible for combating DTT toxicity.

### DTT causes ER stress in part by modulation of the methionine-homocysteine cycle

Disturbance in the methionine-homocysteine cycle (accumulation of homocysteine and depletion of SAM) is known to result in ER stress ^29,33,34^. Since DTT causes ER stress, we asked what the contribution of methionine-homocysteine cycle perturbation in the DTT-induced ER stress was. To this end, we studied the effect of vitamin B12 and methionine supplementation on DTT-induced upregulation of *hsp-4p::GFP*. Supplementation of vitamin B12 and methionine resulted in partial suppression of DTT-induced upregulation of *hsp-4p::GFP* (Figure 5A). Choline supplementation is known to suppress the ER stress caused by the depletion of SAM ^33^. We observed that supplementation of choline resulted in partial suppression of DTT-induced upregulation of *hsp-4p::GFP* (Figure 5A). Thus, these results suggested that DTT causes ER stress in part by modulation of the methionine-homocysteine cycle.

**Figure 5.**
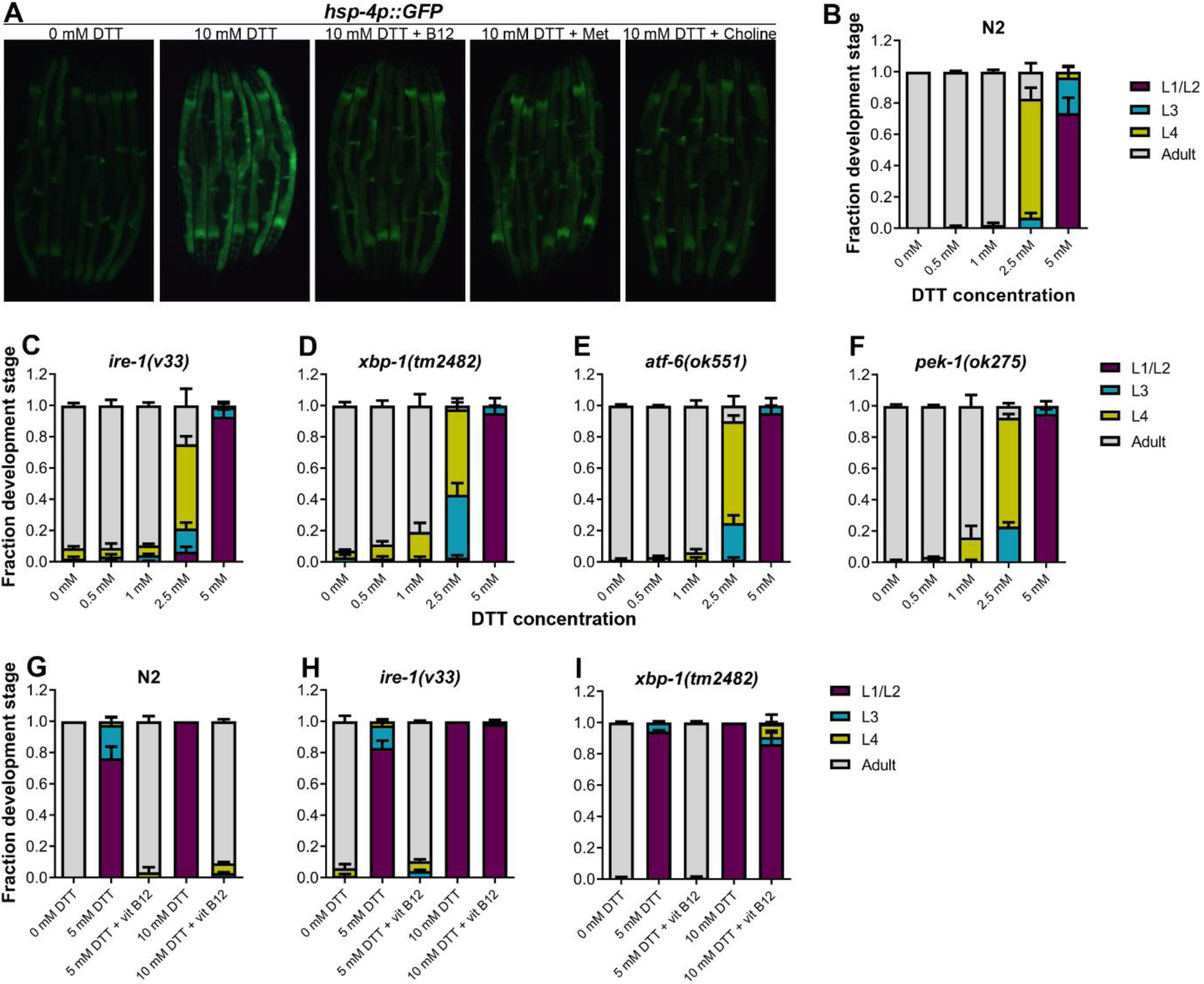
High levels of DTT cause toxicity via the methionine-homocysteine cycle and ER proteotoxic stress. (A) Representative fluorescence images of *hsp-4p::GFP* animals grown on *E. coli* OP50 diet containing 0 mM DTT until the young adult stage, followed by incubation on *E. coli* OP50 diet containing 0 mM DTT, 10 mM DTT, 10 mM DTT supplemented with 50 nM vitamin B12, 10 mM DTT supplemented with 25 mM methionine, and 10 mM DTT supplemented with 80 mM choline for 24 hours. (B-F) Quantification of different developmental stages of N2 (B), *ire-1(v33)* (C), *xbp-1(tm2482)* (D), *atf-6(ok551)* (E), and *pek-1(ok275)* (F) animals on various concentrations of DTT on *E. coli* OP50 diet. The *ire-1(v33)* animals were grown for 84 hours while all other animals were grown for 72 hours at 20°C (n = 3 biological replicates; animals per condition per replicate > 80). (G)-(I) Quantification of development of N2 (G), *ire-1(v33)* (H), and *xbp-1(tm2482)* (I) animals on *E. coli* OP50 plates containing 0 mM DTT, 5 mM DTT, 5 mM DTT supplemented with 50 nM vitamin B12, 10 mM DTT, and 10 mM DTT supplemented with 50 nM vitamin B12 (n = 3 biological replicates; animals per condition per replicate > 80).

### High levels of DTT cause toxicity via the methionine-homocysteine cycle and ER proteotoxic stress

Next, we explored the role of ER stress in DTT-mediated development retardation. The ER UPR pathways alleviate enhanced proteotoxic stress, and therefore, the animals having mutations in the UPR pathways have enhanced development retardation on chemical stressors such as tunicamycin ^35^. Surprisingly, we observed that in the presence of DTT, the mutants of different UPR pathways showed development retardation similar to the N2 animals (Figures 5B-5F). Next, we tested whether vitamin B12 could alleviate DTT toxicity in the different UPR mutants. Similar to the wild-type N2 animals, supplementation of 50 nM vitamin B12 resulted in the rescue of both 5 mM and 10 mM DTT toxicity in *atf-6(ok551)* and *pek-1(ok275)* animals (Figures S6A-S6D). On the other hand, while supplementation of 50 nM vitamin B12 rescued the development of *ire-1(v33)* and *xbp-1(tm2482)* animals on 5 mM DTT, it failed to rescue their development on 10 mM DTT (Figures 5G-5I; Figure S7). Taken together, these results suggested that at lower DTT concentrations, the primary cause of DTT toxicity was the modulation of the methionine-homocysteine cycle, and attenuation of DTT toxicity did not require a functional UPR. On the other hand, at higher DTT concentrations, the toxicity was an outcome of both the modulation of the methionine-homocysteine cycle and the ER proteotoxic stress, and attenuation of the DTT toxicity required a functional IRE-1/XBP-1 UPR pathway.

## DISCUSSION

DTT is a potent reducing agent that effectively reduces protein disulfide bonds ^10,11^, and therefore, has a wide range of applications in protein biochemistry and the pharmaceutical industry. Its antioxidant properties also make it an important candidate drug molecule to treat various pathological conditions associated with oxidative stress ^4,6,8,36^. However, the complete spectrum of physiological effects of DTT inside a cell is not fully understood. The ER is known to be a primary target for DTT inside the cell due to its oxidative nature ^37^. The oxidative nature of the ER is crucial for the formation of disulfide bonds and the proper folding of proteins. DTT leads to the reduction of disulfide bonds in the ER, consequently leading to protein misfolding and ER stress. DTT has been widely used as an ER-specific stressor ^12,13^. Increased ER stress is associated with the activation of cell death programs ^38^, and therefore, the toxicity of DTT has been associated with increased ER stress ^14–16^. In addition to the ER stress, DTT is shown to exert its toxic effects by paradoxically increasing ROS production ^17,18^. In the current study, we showed that in *C. elegans*, DTT causes toxicity by modulating the methionine-homocysteine cycle. By upregulating the expression of SAM-dependent methyltransferase gene *drm-1*, DTT leads to the depletion of SAM that results in toxicity. Supplementation of vitamin B12 and methionine rescues DTT toxicity by repleting SAM levels (Figure 6). Therefore, our study indicates that the effects of DTT are not limited to the ER. Indeed, our data showed that DTT also upregulates the mitochondrial UPR via the methionine-homocysteine cycle and argues against DTT being an ER-specific stressor.

**Figure 6.**
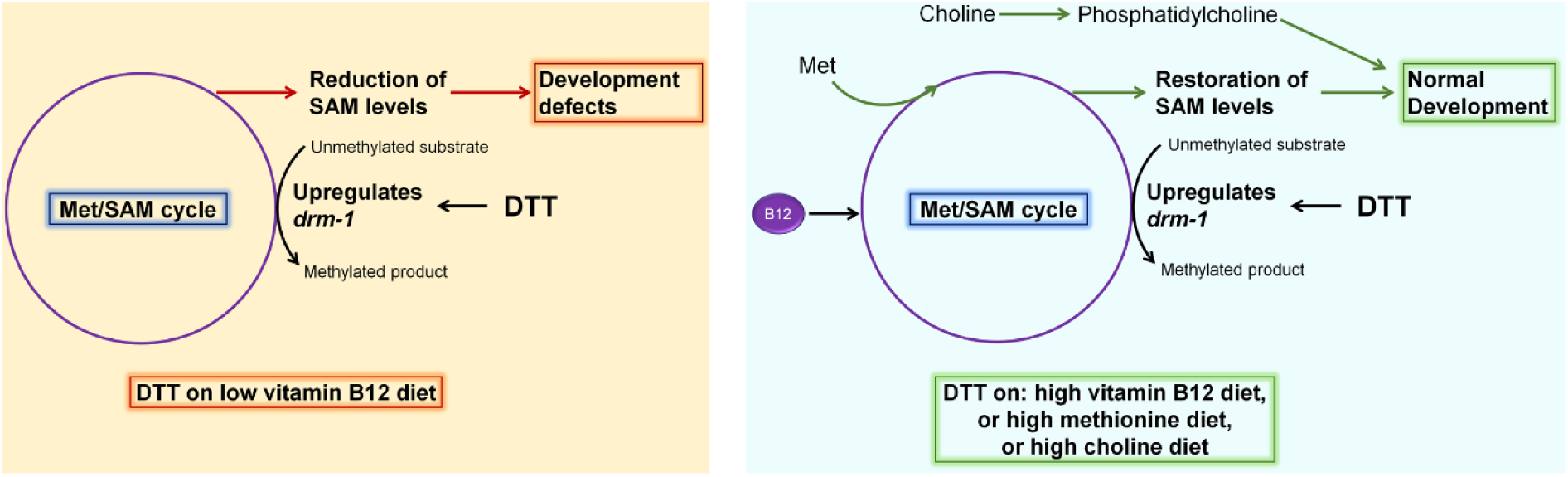
Model for DTT toxicity via the methionine-homocysteine cycle. DTT causes upregulation of the SAM-dependent methyltransferase gene *drm-1*. On low vitamin B12 diet, DTT leads to the depletion of SAM that results in toxicity. Supplementation of vitamin B12 and methionine attenuates DTT toxicity by restoring SAM levels. Supplementation of choline also rescues DTT toxicity by repleting phosphatidylcholine via a SAM-independent pathway.

Our genetic screen for DTT-resistant mutants resulted in the isolation of twelve loss-of-function alleles of the same gene, the SAM-dependent methyltransferase *drm-1*. Mutation in no other gene was recovered, suggesting that the screen had reached a saturation level.

These results indicated that DTT toxicity could be very specifically mediated via the modulation of the methionine-homocysteine cycle. Indeed, we observed that DTT toxicity at 5 mM concentration could be rescued in all of the different ER UPR pathway mutants by supplementation of vitamin B12 (Figure 5; Figures S6 and S7). These results suggested that at 5 mM concentration, the toxic effects of DTT were not due to the ER proteotoxic stress and primarily were due to the modulation of the methionine-homocysteine cycle. However, DTT toxicity at 10 mM DTT concentration could not be rescued by vitamin B12 supplementation in *ire-1(v33)* and *xbp-1(tm2482)* animals, while it was rescued in the wild-type as well as *atf-6(ok551)* and *pek-1(ok275)* animals. This indicated that at 10 mM concentration, DTT toxicity occurred both by the modulation of the methionine-homocysteine cycle and ER proteotoxic stress, and the animals needed a functional IRE-1/XBP-1 pathway to counteract it.

Methionine metabolism and methyltransferases have important roles in regulating health and lifespan ^39^, and defects in the methionine-homocysteine cycle have been linked with a large number of pathological conditions ^40–42^. Therefore, the methionine-homocysteine cycle has become an important intervention point for improving lifespan and health ^39,42–45^. In the current study, we discovered that DTT modulates the methionine-homocysteine cycle resulting in low SAM levels. Reduction of SAM levels is known to enhance the lifespan and metabolic health of a large variety of organisms ^39^. Therefore, it will be intriguing to test whether, at non-toxic concentrations, DTT could have health-improving effects via the methionine-homocysteine cycle.

In the future, it will be interesting to study whether the modulation of the methionine-homocysteine cycle by DTT is a conserved feature across different organisms. In the fungus *Aspergillus niger*, DTT is known to strongly upregulate the expression of two SAM-dependent methyltransferases ^20^. Therefore, it is likely possible that DTT modulates the methionine-homocysteine cycle in *Aspergillus niger*. Similarly, DTT results in increased levels of cysteine in Chinese hamster V79 cells ^46^. Cysteine is synthesized from homocysteine by the transsulfuration pathway, and increased amounts of homocysteine are known to increase cysteine levels ^47^. Therefore, it is possible that DTT modulates the methionine-homocysteine cycle in Chinese hamster V79 cells resulting in increased levels of homocysteine and, consequently, increased cysteine levels. Overall, these studies suggest that modulation of the methionine-homocysteine cycle by DTT could be a conserved feature across different organisms. Future research deciphering the mechanism and relevance of activation of SAM-dependent methyltransferases by DTT will aid in better understanding the physiological effects of DTT.

## MATERIALS AND METHODS

### Bacterial strains

The following bacterial strains were used: *Escherichia coli* OP50, *E. coli* HT115(DE3), *Pseudomonas aeruginosa* PA14 *gacA* mutant, *Comamonas aquatica* DA1877, *Serratia marcescens* Db11, and *Salmonella enterica* serovar Typhimurium 1344. The cultures of these bacteria were grown in Luria-Bertani (LB) broth at 37°C. The nematode growth medium (NGM) plates were seeded with the different bacterial cultures and incubated at room temperature for at least 2 days before using for experiments.

### *C. elegans* strains and growth conditions

*C. elegans* hermaphrodites were maintained at 20°C on NGM plates seeded with *E. coli* OP50 as the food source (Brenner, 1974) unless otherwise indicated. Bristol N2 was used as the wild-type control unless otherwise indicated. The following strains were used in the study: VL749 *wwIs24 [acdh-1p::GFP + unc-119(+)]*, RB755 *metr-1(ok521)*, RB1434 *mmcm-1(ok1637)*, VC2428 *sams-1(ok2946)*, RE666 *ire-1(v33), xbp-1(tm2482)*, RB772 *atf-6(ok551)*, RB545 *pek-1(ok275)*, SJ4100 *zcIs13 [hsp-6p::GFP + lin-15(+)]*, SJ4005 *zcIs4 [hsp-4::GFP]*, and VC20288 *drm-1(gk231506)*. Some of the strains were obtained from the Caenorhabditis Genetics Center (University of Minnesota, Minneapolis, MN). The *xbp-1(tm2482)* animals were obtained from National Bioresource Project, Japan.

### Preparation of NGM plates with different supplements

All the supplements, including DTT, vitamin B12 (cyanocobalamin), methionine, and choline chloride, were dissolved in water to prepare a stock solution. The supplements were added to the nematode growth media just before pouring into plates to obtain a desired final concentration.

### *C. elegans* development assays

Synchronized *C. elegans* eggs were obtained by transferring 15-20 gravid adult hermaphrodites on NGM plates for egg-laying for 2 hours. After 2 hours, the gravid adults were removed. The synchronized eggs were incubated at 20°C for 72 hours. After that, the animals in different development stages (L1/L2, L3, L4, and adult) were quantified. Since *ire-1(v33)* animals develop slightly slower than wild-type animals, the development stages in *ire-1(v33)* animals were quantified after 84 hours of incubation at 20°C. For choline and methionine supplementation assays, the synchronized eggs were incubated at 20°C for 4 days and 5 days, respectively, before quantifying the development stages. Representative images of the NGM plates at the time of quantification of development were also captured. At least three independent experiments (biological replicates) were performed for each condition.

### Rescue of DTT-mediated development retardation by vitamin B12 supplementation

To study whether the DTT-mediated development retardation could be reversed by supplementation of vitamin B12, 2 μL of 750 μM vitamin B12 (total 1.5 nanomoles) was added to the center of a plate with L1-arrested animals after 72 hours of hatching on *E. coli* OP50 with 10 mM DTT. To the control plate with L1-arrested animals after 72 hours of hatching on *E. coli* OP50 with 10 mM DTT, 2 μL of water was added to the center. After that, the plates were incubated at 20°C for 48 hours, followed by the quantification of development stages.

### Forward genetic screen for DTT-resistant mutants

An ethyl methanesulfonate (EMS) mutagenesis selection screen ^48^ was performed using the wild-type N2 strain. Approximately 2500 synchronized late L4 larvae were treated with 50 mM EMS for 4 hours and then washed three times with M9 medium. The washed animals (P0 generation) were then transferred to 9-cm NGM plates containing *E. coli* OP50 and allowed to lay eggs (F1 progeny) overnight. The P0s were then washed away with M9 medium, while the F1 eggs remained attached to the bacterial lawn. The F1 eggs were allowed to grow to adulthood. The adult F1 animals were bleached to obtain F2 eggs. The F2 eggs were transferred to *E. coli* OP50 plates containing 10 mM DTT and incubated at 20°C for 72 hours. After that, the plates were screened for animals that developed to L4 or adult stages. Approximately 50,000 haploid genomes were screened, and twelve mutants were isolated. All of the mutants were backcrossed six times with the parental N2 strain before analysis.

### Whole-genome sequencing (WGS) and data analysis

The genomic DNA was isolated as described earlier ^49^. Briefly, the mutant animals were grown at 20°C on NGM plates seeded with *E. coli* OP50 until starvation. The animals were rinsed off the plates with M9, washed three times, incubated in M9 with rotation for 2 hours to eliminate food from the intestine, and washed three times again with M9. Genomic DNA extraction was performed using the Gentra Puregene Kit (Qiagen, Netherlands). DNA libraries were prepared according to a standard Illumina (San Diego, CA) protocol. The DNA was subjected to WGS on an Illumina HiSeq sequencing platform using 150 paired-end nucleotide reads. Library preparation and WGS were performed at Clevergene Biocorp Pvt. Ltd, Bengaluru, India.

The whole-genome sequence data were analyzed using the web platform Galaxy. The forward and reverse FASTQ reads, *C. elegans* reference genome Fasta file (ce11M.fa), and the gene annotation file (SnpEff4.3 WBcel235.86) were input into the Galaxy workflow. The low-quality ends of the FASTQ reads were trimmed using the Sickle tool. The trimmed FASTQ reads were mapped to the reference genome Fasta files with the BWA-MEM tool. Using the MarkDuplicates tool, any duplicate reads (mapped to multiple sites) were filtered. Subsequently, the variants were detected using the FreeBayes tool that finds small polymorphisms, including single-nucleotide polymorphisms (SNPs), insertions and deletions (indels), multi-nucleotide polymorphisms (MNPs), and complex events (composite insertion and substitution events) smaller than the length of a short-read sequencing alignment. The common variants among different mutants were subtracted. The SnpEff4.3 WBcel235.86 gene annotation file was used to annotate and predict the effects of genetic variants (such as amino acid changes). Finally, the linkage maps for each mutant were generated using the obtained variation.

### Plasmid constructs and generation of transgenic *C. elegans*

For rescue of the DTT-resistance phenotype of *drm-1(jsn11)* animals, the *drm-1* gene along with its promoter region (918-bp upstream) was amplified from genomic DNA of N2 animals. The gene, including its stop codon, was cloned in the pPD95.77 plasmid containing the SL2 sequence before GFP. For generating the *drm-1p::GFP* reporter strain, the promoter region of the *drm-1* gene (918-bp upstream) was amplified from genomic DNA of N2 animals and cloned in the pPD95.77 plasmid before GFP. *drm-1(jsn11)* animals were microinjected with *drm-1p::drm-1::SL2::GFP* plasmid along with pCFJ90 (*myo-2p::mCherry*) as a coinjection marker to generate rescue strains. N2 wild-type animals were microinjected with *drm-1p::GFP* plasmid along with pCFJ90 as a coinjection marker to generate *drm-1p::GFP* reporter strain. The *drm-1p::drm-1::SL2::GFP* and *drm-1p::GFP* plasmids were used at a concentration of 50 ng/μl, while the coinjection marker was used at a concentration of 5 ng/μl. The plasmids were maintained as extrachromosomal arrays, and at least two independent lines were maintained for each plasmid.

### RNA isolation and quantitative reverse transcription-PCR (qRT-PCR)

Animals were synchronized by egg laying. Approximately 35 N2 gravid adult animals were transferred to 9-cm *E. coli* OP50 plates without DTT and allowed to lay eggs for 4 hours. The gravid adults were then removed, and the eggs were allowed to develop at 20°C for 72 hours. Subsequently, the synchronized adult animals were collected with M9, washed twice with M9, and then transferred to 9-cm *E. coli* OP50 plates containing 10 mM DTT. The control animals were maintained on *E. coli* OP50 plates without DTT. After the transfer of the animals, the plates were incubated at 20°C for 4 hours. Subsequently, the animals were collected, washed with M9 buffer, and frozen in TRIzol reagent (Life Technologies, Carlsbad, CA). Total RNA was extracted using the RNeasy Plus Universal Kit (Qiagen, Netherlands). A total of 1 μg of total RNA was reverse-transcribed with oligo dT primers using the Transcriptor High Fidelity cDNA Synthesis Kit (Roche) according to the manufacturer’s protocols. qRT-PCR was conducted using TB Green fluorescence (TaKaRa) on a LightCycler 480 II System (Roche Diagnostics) in 96-well-plate format. Twenty microliter reactions were analyzed as outlined by the manufacturer (TaKaRa). The relative fold-changes of the transcripts were calculated using the comparative *CT*(2^-ΔΔ*CT*^) method and normalized to pan-actin (*act-1, -3, -4*) as described earlier ^50^. All samples were run in triplicate.

### *drm-1p::GFP* induction by DTT

The *drm-1p::GFP* animals were synchronized by egg laying on *E. coli* OP50 plates without DTT and incubated at 20°C for 72 hours. Subsequently, the animals were transferred to *E. coli* OP50 plates containing 10 mM DTT and incubated at 20°C for 8 hours. The control animals were maintained on *E. coli* OP50 plates without DTT. After that, the animals were prepared for fluorescence imaging.

### *acdh-1p::GFP* assays

The *acdh-1p::GFP* animals were synchronized by egg laying on different bacterial diets without DTT. The animals were incubated at 20°C for 72 hours. After that, the animals were prepared for fluorescence imaging.

### *hsp-6p::GFP* induction assays

The *hsp-6p::GFP* animals were synchronized by egg laying on *E. coli* OP50 plates without DTT and incubated at 20°C for 72 hours. Subsequently, the animals were transferred to *E. coli* OP50 plates containing 0 mM DTT, 10 mM DTT, and 10 mM DTT supplemented with 50 nM vitamin B12 followed by incubation at 20°C for 24 hours. After that, the animals were prepared for fluorescence imaging.

### *hsp-4p::GFP* induction assays

The *hsp-4p::GFP* animals were synchronized by egg laying on *E. coli* OP50 plates without DTT and incubated at 20°C for 72 hours. Subsequently, the animals were transferred to *E. coli* OP50 plates containing 0 mM DTT, 10 mM DTT, 10 mM DTT supplemented with 50 nM vitamin B12, 10 mM DTT supplemented with 25 mM methionine, and 10 mM DTT supplemented with 80 mM choline chloride followed by incubation at 20°C for 24 hours. After that, the animals were prepared for fluorescence imaging.

### Fluorescence imaging

Fluorescence imaging was carried out as described previously ^51^. Briefly, the animals were anesthetized using an M9 salt solution containing 50 mM sodium azide and mounted onto 2% agar pads. The animals were then visualized using a Nikon SMZ-1000 fluorescence stereomicroscope.

### Quantification and Statistical Analysis

The statistical analysis was performed with Prism 8 (GraphPad). All error bars represent the standard deviation (SD). The two-sample t test was used when needed, and the data were judged to be statistically significant when p < 0.05. In the figures, asterisks (*) denote statistical significance as follows: *, p < 0.05, **, p < 0.01, ***, p < 0.001, as compared with the appropriate controls.

## SUPPLEMENTAL INFORMATION

The Supplemental Information includes seven figures and one table.

## ACKNOWLEDGMENTS

This work was supported by IISER Bhopal (Grant # INST/BIO/2019091), IISER Mohali intramural funds, Science and Engineering Research Board (SERB) Startup Research Grant (Ref. No. SRG/2020/000022) awarded by DST, India, and Ramalingaswami Re-entry Fellowship (Ref. No. BT/RLF/Re-entry/50/2020) awarded by the Department of Biotechnology, India. G.G. was supported by an INSPIRE-SHE Scholarship from DST, India. Some strains used in this study were provided by the Caenorhabditis Genetics Center (CGC), which is funded by the NIH Office of Research Infrastructure Programs (P40 OD010440). We thank Dr. Pratima Pandey (IISER Mohali) for help with qRT-PCR experiments, Ravi (IISER Mohali) for some technical assistance, and Dr. Lolitika Mandal’s laboratory for *acdh-1p::GFP* fluorescence imaging.

## AUTHOR CONTRIBUTIONS

G.G., and J.S. conceived and designed the experiments. G.G., and J.S. performed the experiments. J.S. analyzed the data and wrote the paper.

## DATA AVAILABILITY

The whole-genome sequence data for JSJ1-JSJ12 have been submitted to the public repository, the Sequence Read Archive, with BioProject ID PRJNA786771.

## Supplementary Figures

**Figure S1.**
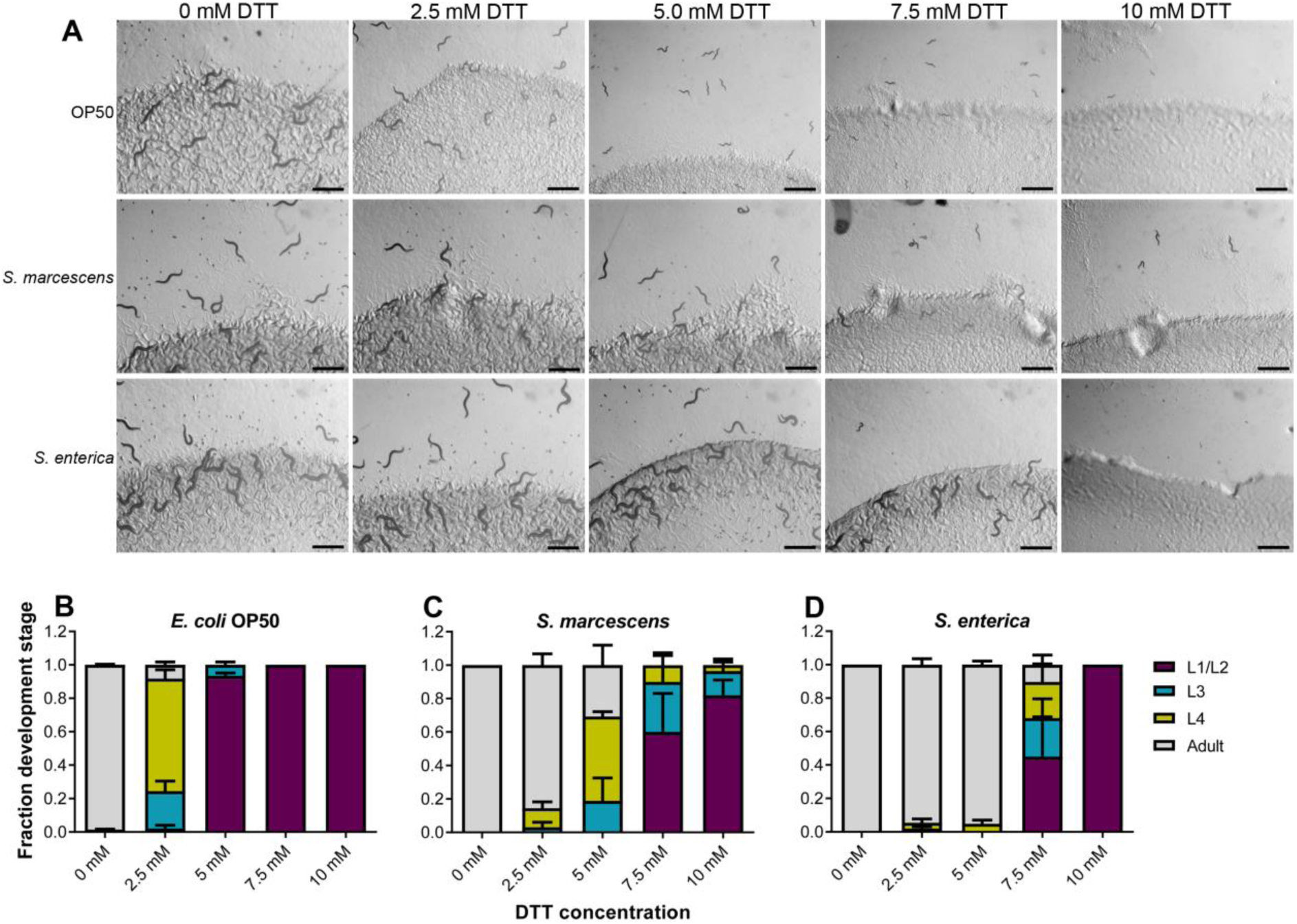
DTT affects *C. elegans* development in a diet-dependent manner. (A) Representative images of wild-type N2 *C. elegans* on various concentrations of DTT on *E. coli* OP50, *S. marcescens* Db11, and *S. enterica* diets after 72 hours of hatching at 20°C. Scale bar = 1 mm. (B-D) Quantification of different developmental stages of wild-type N2 *C. elegans* on various concentrations of DTT on *E. coli* OP50 (B), *S. marcescens* Db11 (C), and *S. enterica* (D) diets after 72 hours of hatching at 20°C (n = 3 biological replicates; animals per condition per replicate > 80).

**Figure S2.**
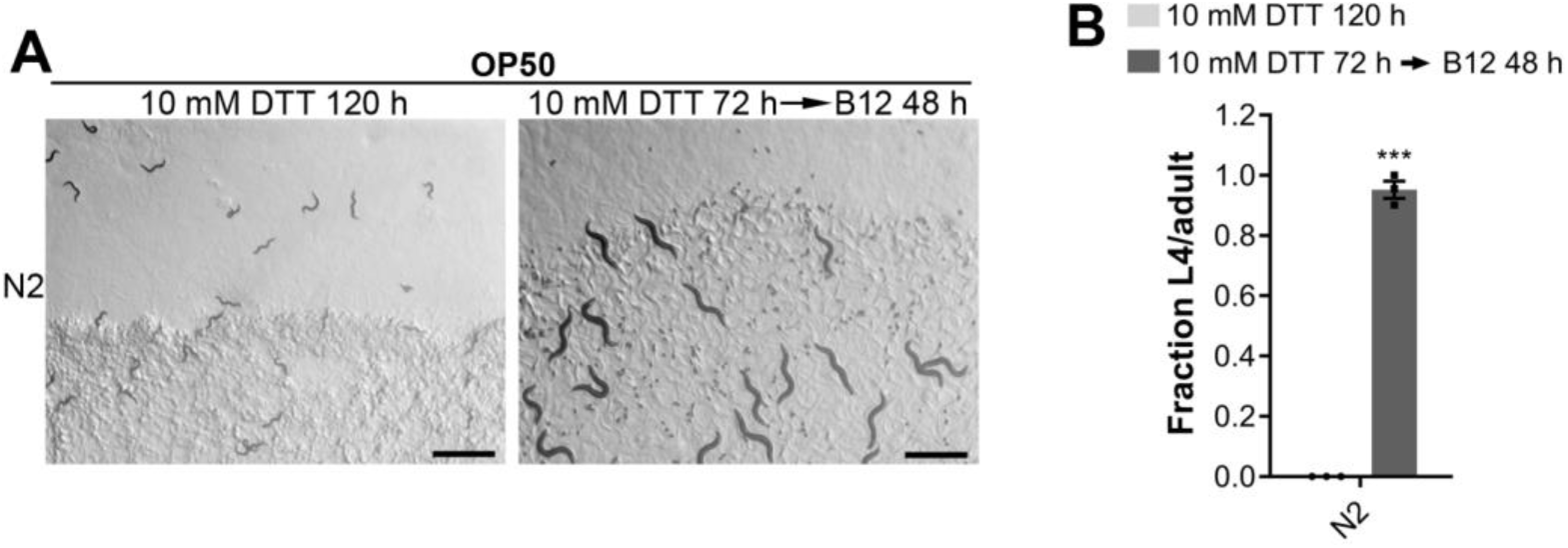
Vitamin B12 supplementation reverses DTT toxicity. (A) Representative images of wild-type N2 *C. elegans* grown on *E. coli* OP50 diet containing 10 mM DTT for 72 hours, followed by another 48 hours growth upon supplementation of either water blank (left panel) or 1.5 nanomoles of vitamin B12 (right panel). Scale bar = 1 mm. (B) Fraction L4 or adult wild-type N2 *C. elegans* grown on *E. coli* OP50 diet containing 10 mM DTT for 72 hours, followed by another 48 hours growth upon supplementation of either water blank (10 mM DTT 120 h) or 1.5 nanomoles of vitamin B12. ***p < 0.001 via the t test (n = 3 biological replicates; animals per condition per replicate > 60).

**Figure S3.**
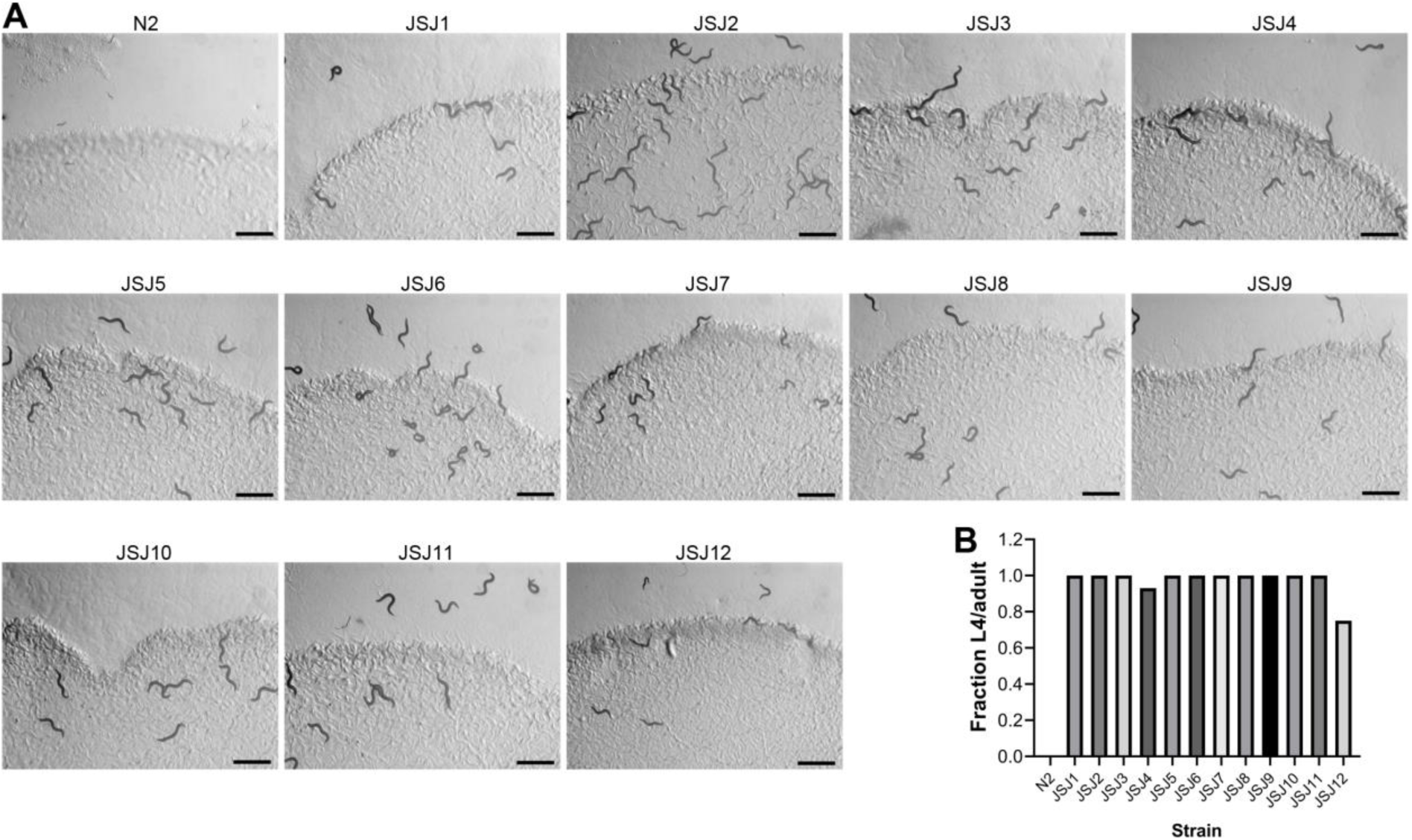
Forward genetic screen resulted in the isolation of twelve DTT-resistant mutants. (A) Representative images of N2 and DTT-resistant mutants (JSJ1 to JSJ12) on *E. coli* OP50 diet containing 10 mM DTT after 72 hours of hatching at 20°C. Scale bar = 1 mm. (B) Fraction L4 or adult N2 and DTT-resistant mutants on *E. coli* OP50 diet containing 10 mM DTT after 72 hours of hatching at 20°C.

**Figure S4.**
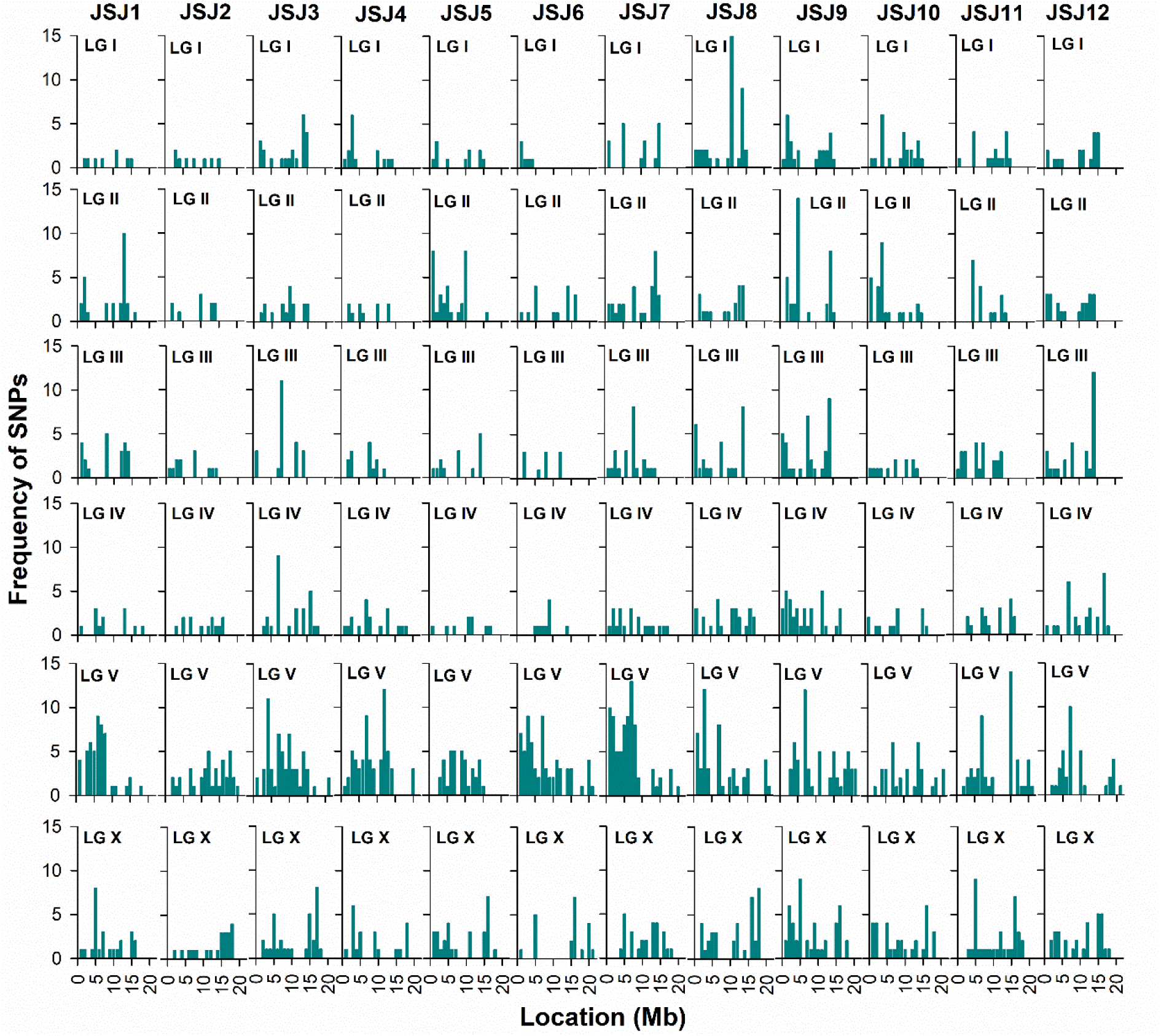
Mapping of the mutations by whole-genome sequencing. The frequencies of single-nucleotide polymorphisms (SNPs) were plotted against the positions of different chromosomes or linkage groups (LG). The high frequency of SNPs on a particular chromosome compared to other regions of the genome represents the region linked with the causative mutation(s). All mutants showed a high frequency of SNPs on chromosome V.

**Figure S5.**
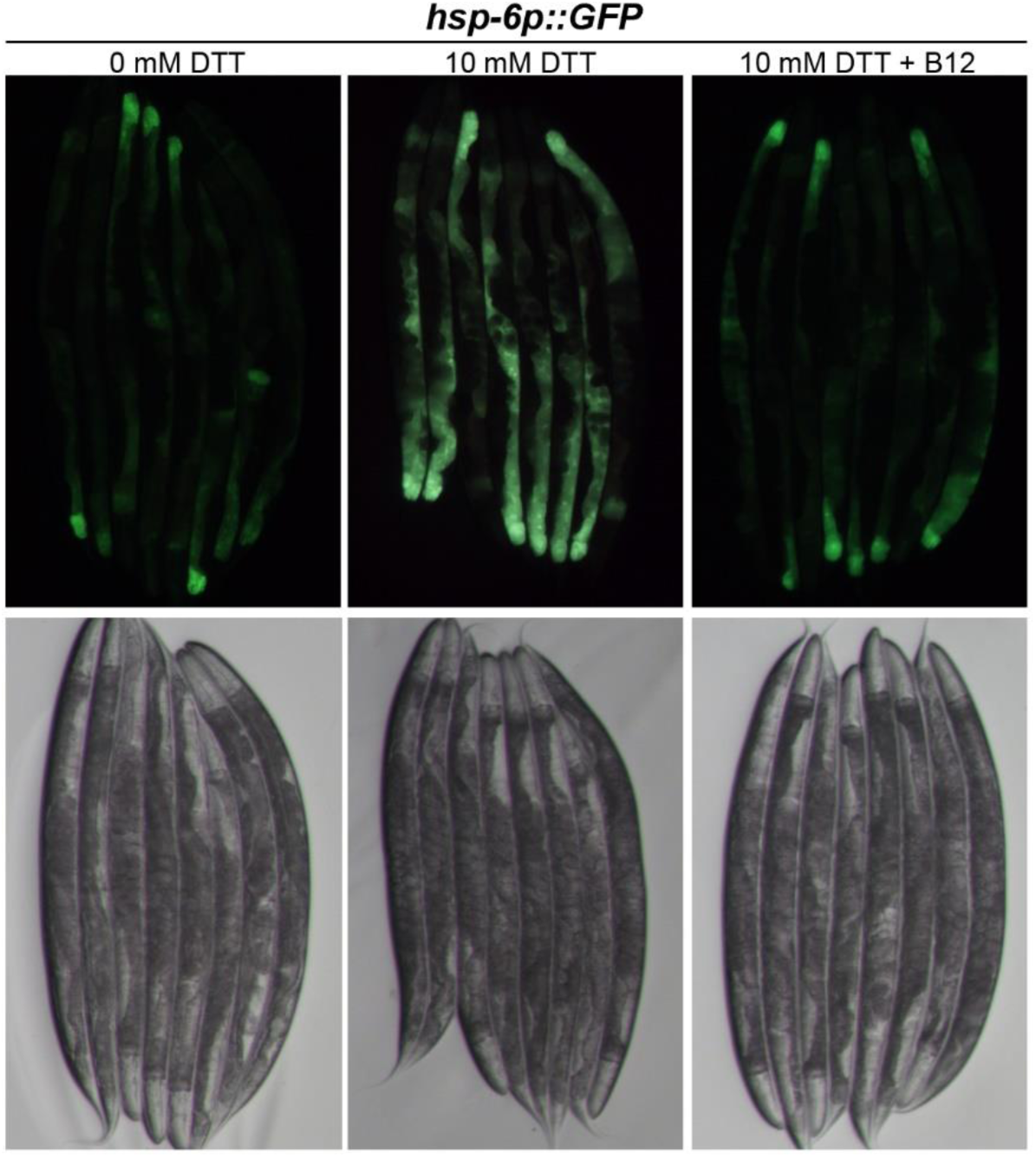
DTT upregulates the mitochondrial UPR. Representative fluorescence images (top) and the corresponding brightfield images (bottom) of *hsp-6p::GFP* animals grown on *E. coli* OP50 diet containing 0 mM DTT until the young adult stage, followed by incubation on *E. coli* OP50 diet containing 0 mM DTT, 10 mM DTT, and 10 mM DTT supplemented with 50 nM vitamin B12 for 24 hours.

**Figure S6.**
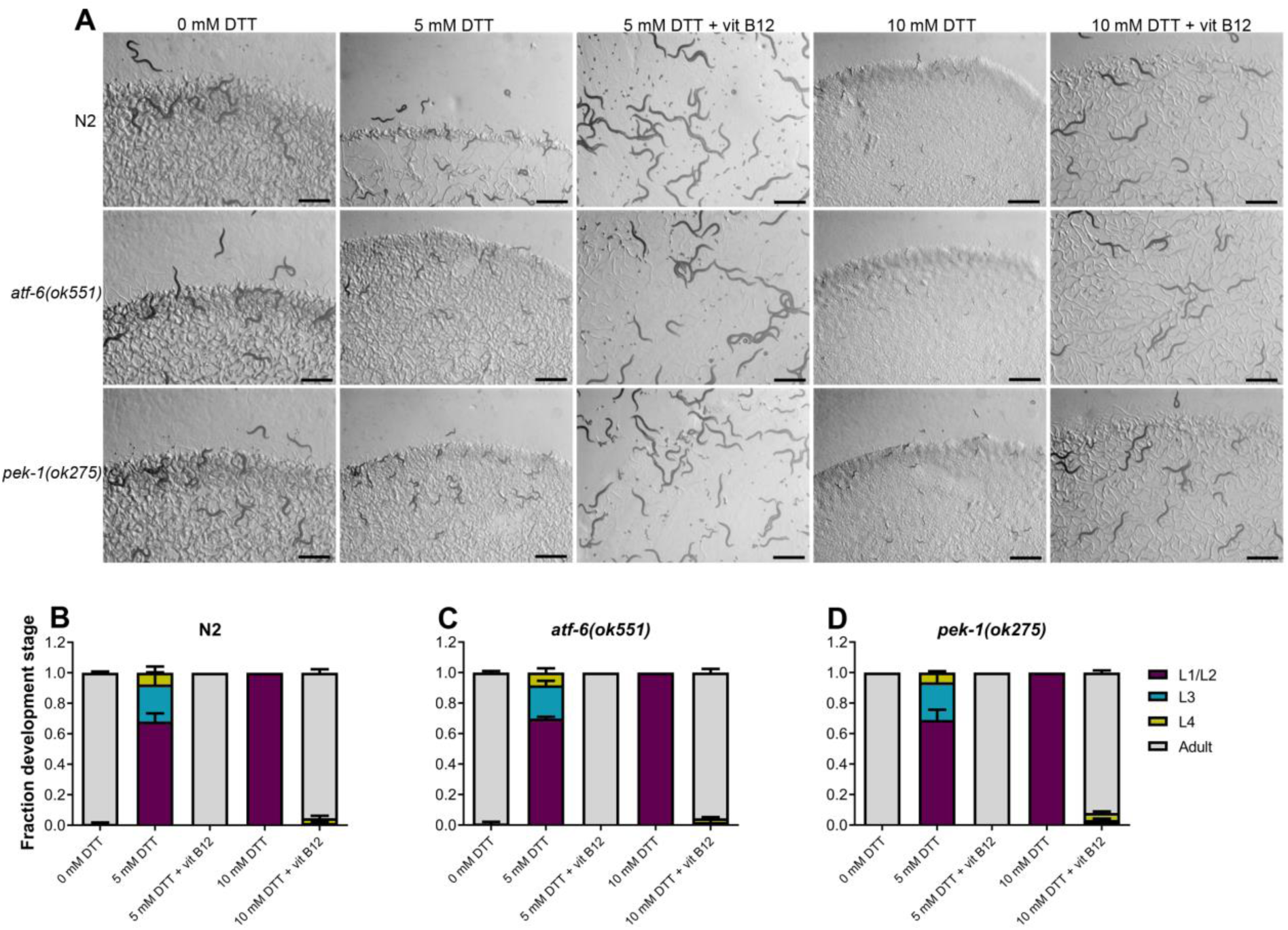
The ATF-6 and PEK-1 pathways are not involved in combating DTT toxicity. (A) Representative images of N2, *atf-6(ok551)*, and *pek-1(ok275)* animals on *E. coli* OP50 plates containing 0 mM DTT, 5 mM DTT, 5 mM DTT supplemented with 50 nM vitamin B12, 10 mM DTT, and 10 mM DTT supplemented with 50 nM vitamin B12. Scale bar = 1 mm. (B-D) Quantification of different developmental stages of N2 (B), *atf-6(ok551)* (C), and *pek-1(ok275)* (D) animals on *E. coli* OP50 plates containing 0 mM DTT, 5 mM DTT, 5 mM DTT supplemented with 50 nM vitamin B12, 10 mM DTT, and 10 mM DTT supplemented with 50 nM vitamin B12 (n = 3 biological replicates; animals per condition per replicate > 80).

**Figure S7.**
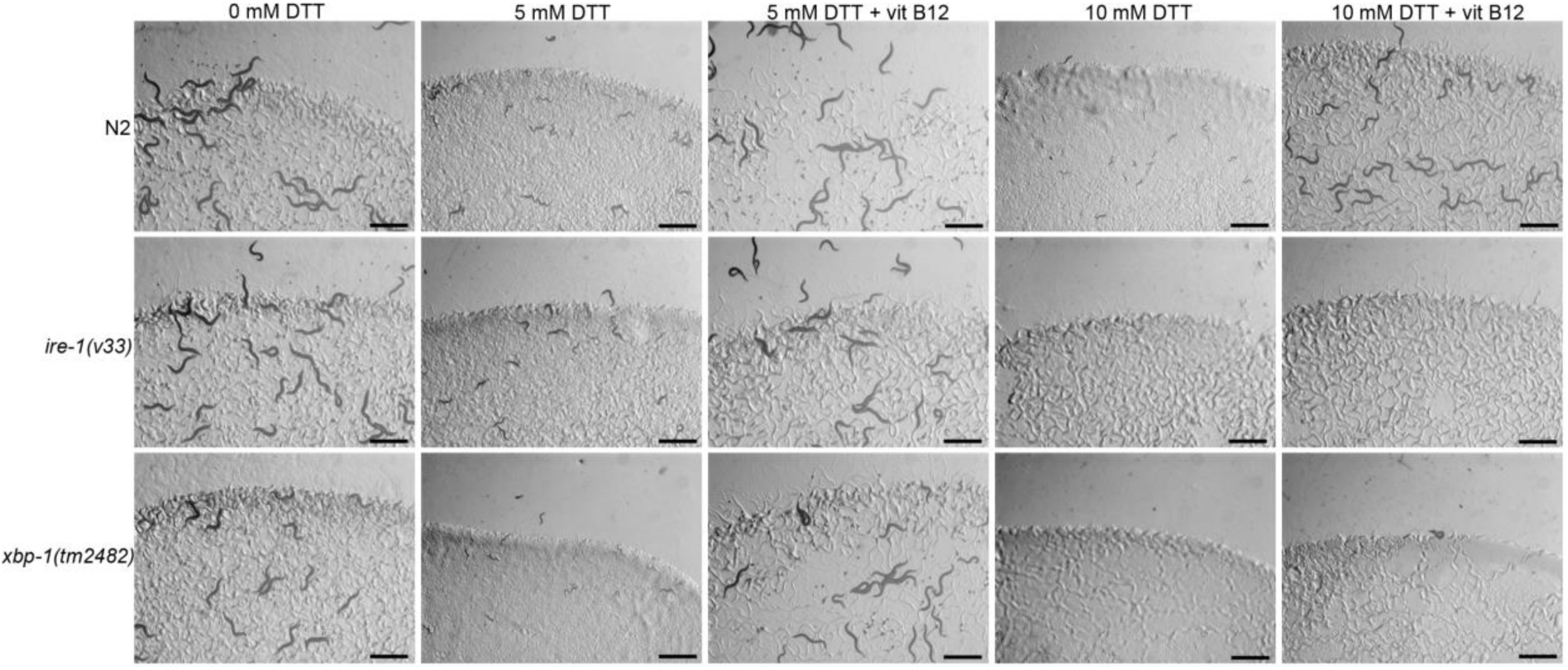
Counteracting high, but not low, levels of DTT requires a functional IRE-1/XBP-1 UPR pathway. Representative images of N2, *ire-1(v33)*, and *xbp-1(tm2482)* animals on *E. coli* OP50 plates containing 0 mM DTT, 5 mM DTT, 5 mM DTT supplemented with 50 nM vitamin B12, 10 mM DTT, and 10 mM DTT supplemented with 50 nM vitamin B12. Scale bar = 1 mm.

**Table S1:**
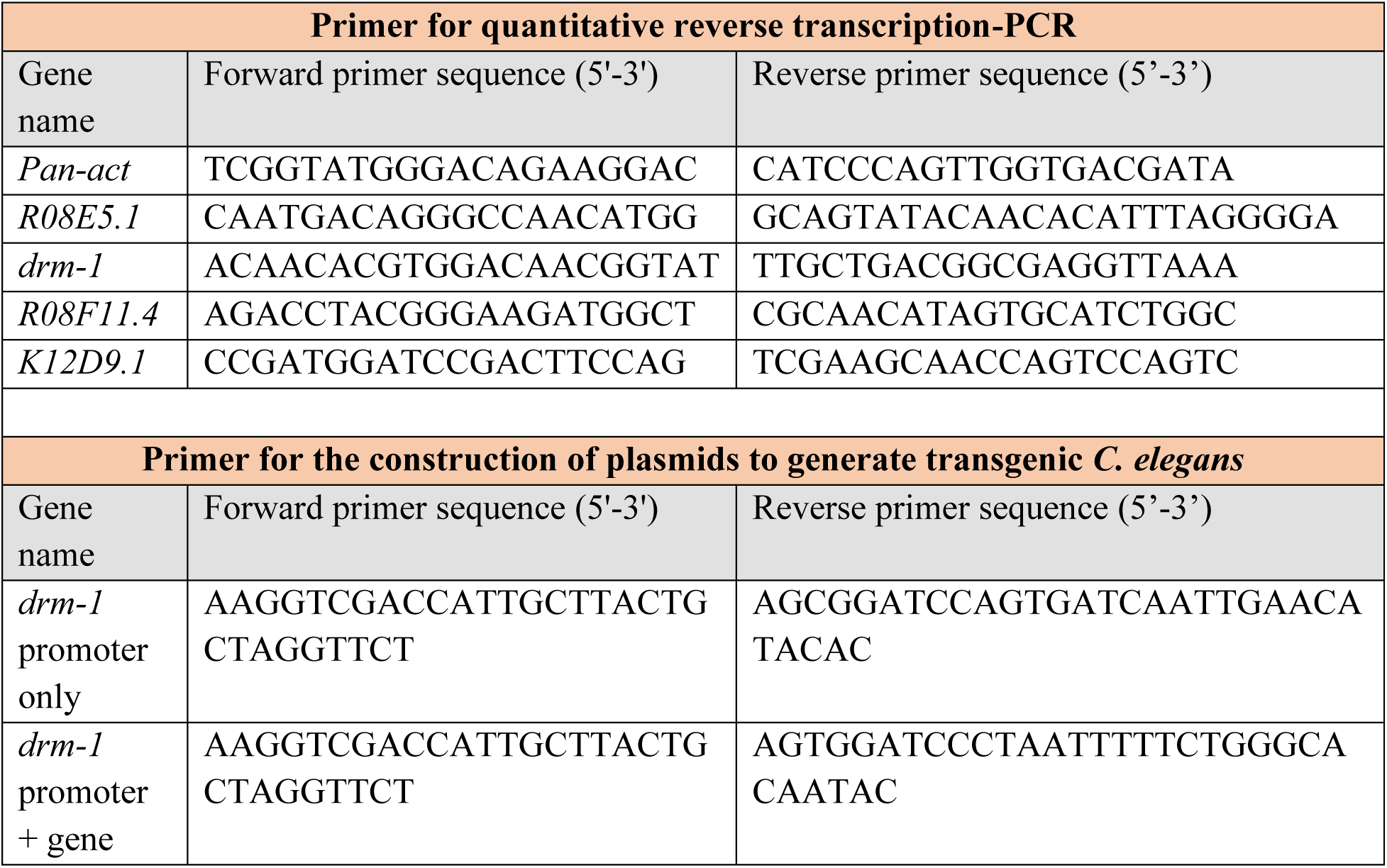
Primers used in the study.

## REFERENCES

1. Trachootham, D., Lu, W., Ogasawara, M. A., Valle, N.R. Del & Huang, P. Redox regulation of cell survival. Antioxidants Redox Signal. 10, 1343–1374 (2008).

2. Sies, H. & Jones, D. P. Reactive oxygen species (ROS) as pleiotropic physiological signalling agents. Nat. Rev. Mol. Cell Biol. 21, 363–383 (2020).

3. Kurutas, E. B. The importance of antioxidants which play the role in cellular response against oxidative/nitrosative stress : current state. Nutr. J. 15, 71 (2016).

4. Heidari, R. et al. Dithiothreitol supplementation mitigates hepatic and renal injury in bile duct ligated mice: Potential application in the treatment of cholestasis-associated complications. Biomed. Pharmacother. 99, 1022–1032 (2018).

5. Heidari, R. et al. Sulfasalazine-induced renal injury in rats and the protective role of thiol-reductants. Ren. Fail. 38, 137–141 (2016).

6. Li, K., Zhang, J., Cao, J., Li, X. & Tian, H. 1,4-Dithiothreitol treatment ameliorates hematopoietic and intestinal injury in irradiated mice: Potential application of a treatment for acute radiation syndrome. Int. Immunopharmacol. 76, 105913 (2019).

7. Pocernich, C. B. & Butterfield, D. A. Elevation of glutathione as a therapeutic strategy in Alzheimer disease. Biochim. Biophys. Acta - Mol. Basis Dis. 1822, 625–630 (2012).

8. Offen, D., Ziv, I., Sternin, H., Melamed, E. & Hochman, A. Prevention of Dopamine-Induced Cell Death by Thiol Antioxidants: Possible Implications for Treatment of Parkinson’s Disease. Exp. Neurol. 39, 32–39 (1996).

9. Gusarov, I. et al. Dietary thiols accelerate aging of C. elegans. Nat. Commun. 12, 4336 (2021).

10. Cleland, W. W. Dithiothreitol, a New Protective Reagent for SH Groups. Biochemistry 3, 480–482 (1964).

11. Konigsberg, W. Reduction of Disulfide Bonds in Proteins with Dithiothreitol. Methods Enzymol. 25, 185–188 (1972).

12. Braakman, I., Helenius, J. & Helenius, A. Manipulating disulfide bond formation and protein folding in the endoplasmic reticulum. EMBO J. 11, 1717–1722 (1992).

13. Oslowski, C. M. & Urano, F. Measuring ER Stress and the Unfolded Protein Response Using Mammalian Tissue Culture System. Methods Enzymol. 490, 71–92 (2011).

14. Li, B. et al. Differences in endoplasmic reticulum stress signalling kinetics determine cell survival outcome through activation of MKP-1. Cell. Signal. 23, 35–45 (2011).

15. Xiang, X. Y. et al. Inhibition of autophagic flux by ROS promotes apoptosis during DTT-induced ER/oxidative stress in HeLa cells. Oncol. Rep. 35, 3471–3479 (2016).

16. Qin, L., Wang, Z., Tao, L. & Wang, Y. ER stress negatively regulates AKT/TSC/mTOR pathway to enhance autophagy. Autophagy 6, 239–247 (2010).

17. Held, K. D., Sylvester, F. C., Hopcia, K. L. & Biaglow, J. E. Role of Fenton Chemistry in Thiol-Induced Toxicity and Apoptosis. Radiat. Res. 145, 542–553 (1996).

18. Held, K. D. & Melder, D. C. Toxicity of the Sulfhydryl-Containing Radioprotector Dithiothreitol. Radiat. Res. 112, 544–554 (1987).

19. Guillemette, T. et al. Genomic analysis of the secretion stress response in the enzyme-producing cell factory Aspergillus niger. BMC Genomics 8, 158 (2007).

20. MacKenzie, D. A. et al. UPR-independent dithiothreitol stress-induced genes in Aspergillus niger. Mol. Genet. Genomics 274, 410–418 (2005).

21. Tartier, L., McCarey, Y. L., Biaglow, J. E., Kochevar, I. E. & Held, K. D. Apoptosis induced by dithiothreitol in HL-60 cells shows early activation of caspase 3 and is independent of mitochondria. Cell Death Differ. 7, 1002–1010 (2000).

22. Messias Sandes, J. et al. The effects of endoplasmic reticulum stressors, tunicamycin and dithiothreitol on Trypanosoma cruzi. Exp. Cell Res. 383, 111560 (2019).

23. Kozlowski, L., Garvis, S., Bedet, C. & Palladino, F. The Caenorhabditis elegans HP1 family protein HPL-2 maintains ER homeostasis through the UPR and hormesis. Proc. Natl. Acad. Sci. 111, 5956–5961 (2014).

24. Tan, M. W., Mahajan-Miklos, S. & Ausubel, F. M. Killing of Caenorhabditis elegans by Pseudomonas aeruginosa used to model mammalian bacterial pathogenesis. Proc. Natl. Acad. Sci. U. S. A. 96, 715–20 (1999).

25. Watson, E. et al. Interspecies systems biology uncovers metabolites affecting C. elegans gene expression and life history traits. Cell 156, 759–770 (2014).

26. Revtovich, A. V., Lee, R. & Kirienko, N. V. Interplay between mitochondria and diet mediates pathogen and stress resistance in caenorhabditis elegans. PLoS Genet. 15, e1008011 (2019).

27. Wei, W. & Ruvkun, G. Lysosomal activity regulates Caenorhabditis elegans mitochondrial dynamics through vitamin B12 metabolism. Proc. Natl. Acad. Sci. U. S. A. 117, 19970–19981 (2020).

28. Amin, M. R., Mahmud, S. A., Dowgielewicz, J. L., Sapkota, M. & Pellegrino, M. W. A novel gene-diet interaction promotes organismal lifespan and host protection during infection via the mitochondrial UPR. PLoS Genet. 16, e1009234 (2020).

29. Walker, A. K. et al. A conserved SREBP-1/phosphatidylcholine feedback circuit regulates lipogenesis in metazoans. Cell 147, 840–852 (2011).

30. Vance, J. E. & Vance, D. E. Phospholipid biosynthesis in mammalian cells. Biochem. Cell Biol. 82, 113–128 (2004).

31. Giese, G. E. et al. Caenorhabditis elegans methionine/S-adenosylmethionine cycle activity is sensed and adjusted by a nuclear hormone receptor. Elife 9, e60259 (2020).

32. Ding, W. et al. S-adenosylmethionine levels govern innate immunity through distinct methylation-dependent pathways. Cell Metab. 22, 633–645 (2015).

33. Hou, N. S. et al. Activation of the endoplasmic reticulum unfolded protein response by lipid disequilibrium without disturbed proteostasis in vivo. Proc. Natl. Acad. Sci. 111, E2271–E2280 (2014).

34. Werstuck, G. H. et al. Homocysteine-induced endoplasmic reticulum stress causes dysregulation of the cholesterol and triglyceride biosynthetic pathways. J. Clin. Invest. 107, 1263–1273 (2001).

35. Shen, X. et al. Complementary Signaling Pathways Regulate the Unfolded Protein Response and Are Required for C. elegans Develoment. Cell 107, 893–903 (2001).

36. Heidari, R., Esmailie, N., Azarpira, N., Najibi, A. & Niknahad, H. Effect of thiol-reducing agents and antioxidants on sulfasalazine-induced hepatic injury in normotermic recirculating isolated perfused rat liver. Toxicol. Res. 32, 133–140 (2016).

37. Merksamer, P. I., Trusina, A. & Papa, F. R. Real-Time Redox Measurements during Endoplasmic Reticulum Stress Reveal Interlinked Protein Folding Functions. Cell 135, 933–947 (2008).

38. Sano, R. & Reed, J. C. ER stress-induced cell death mechanisms. Biochim. Biophys. Acta - Mol. Cell Res. 1833, 3460–3470 (2013).

39. Parkhitko, A. A., Jouandin, P., Mohr, S. E. & Perrimon, N. Methionine metabolism and methyltransferases in the regulation of aging and lifespan extension across species. Aging Cell 18, e13034 (2019).

40. Finkelstein, J. D. Inborn Errors of Sulfur-Containing Amino Acid Metabolism. J. Nutr. 136, 1750S–1754S (2006).

41. Li, Z. et al. Methionine metabolism in chronic liver diseases: an update on molecular mechanism and therapeutic implication. Signal Transduct. Target. Ther. 5, 280 (2020).

42. Sanderson, S. M., Gao, X., Dai, Z. & Locasale, J. W. Methionine metabolism in health and cancer: a nexus of diet and precision medicine. Nat. Rev. Cancer 19, 625–637 (2019).

43. Lam, A. B., Kervin, K. & Tanis, J. E. Vitamin B12 impacts amyloid beta-induced proteotoxicity by regulating the methionine/S-adenosylmethionine cycle. Cell Rep. 36, 109753 (2021).

44. Li, H., Lu, H., Tang, W. & Zuo, J. Targeting methionine cycle as a potential therapeutic strategy for immune disorders. Expert Opin. Ther. Targets 21, 861–877 (2017).

45. Konno, M. et al. The one-carbon metabolism pathway highlights therapeutic targets for gastrointestinal cancer. Int. J. Oncol. 50, 1057–1063 (2017).

46. Dennist, M. F., Stratford, M. R. L., Wardman, P. & Watfa, R. R. Increase in intracellular cysteine after exposure to dithiothreitol: implications in radiobiology. Int. J. Radiat. Biol. 56, 877–883 (1989).

47. Sbodio, J. I., Snyder, S. H. & Paul, B. D. Regulators of the transsulfuration pathway. Br. J. Pharmacol. 176, 583–593 (2019).

48. Singh, J. Harnessing the power of genetics: fast forward genetics in Caenorhabditis elegans. Mol. Genet. Genomics 296, 1–20 (2021).

49. Singh, J. & Aballay, A. Endoplasmic reticulum stress caused by lipoprotein accumulation suppresses immunity against bacterial pathogens and contributes to immunosenescence. MBio 8, e00778–17 (2017).

50. Singh, J. & Aballay, A. Microbial colonization activates an immune fight-and-flight response via neuroendocrine signaling. Dev. Cell 49, 89–99 (2019).

51. Singh, J. & Aballay, A. Intestinal infection regulates behavior and learning via neuroendocrine signaling. Elife 8, e50033 (2019).

